# Genetic structures of the Japanese stone loach *Barbatula oreas* (Cypriniformes: Nemacheilidae) in Sakhalin and Hokkaido: back dispersal from Hokkaido to Sakhalin

**DOI:** 10.64898/2026.06.17.732794

**Authors:** Hayato Niinuma, Kensuke Kobayashi, Masaki Takenaka, Gaku Ueki, Sergei V. Shedko, Tatyana S. Vshivkova, Koji Tojo

## Abstract

Understanding how dispersal and vicariance shape species distributions is a central goal in biogeography, yet the role of islands as sources of continental diversity remains poorly resolved. While multiple dispersal routes from the Eurasian continent to the Japanese Archipelago have been proposed, back dispersal from islands to the mainland is rarely documented, particularly in primary freshwater taxa constrained by marine barriers. Here, we investigated the population genetic structure and phylogeographic history of the Japanese stone loach *Barbatula oreas*, distributed in Hokkaido and Sakhalin, using mtDNA, nDNA, and genome-wide SNP data. We identified two differentiated Northern and Southern lineages within Hokkaido that diverged during the Pleistocene, indicating that geological events such as paleo-catchment reorganization, mountain uplift, and volcanic activity have shaped the present population structure. Ancestral area reconstruction based on mtDNA phylogeny identified Hokkaido as the origin of *B. oreas* and revealed dispersal from Hokkaido to Sakhalin, indicating back dispersal from islands toward the mainland. This pattern contrasts with the prevailing hypothesis of southward colonization from the continent via Sakhalin to Hokkaido. Additionally, low genetic differentiation between Hokkaido and Sakhalin suggested genetic exchange across the strait, consistent with paleo-catchment reconstruction indicating past catchment connectivity between the regions. These results highlight the combined roles of geological dynamics and sea-level fluctuations in shaping genetic structure, challenge the conventional continent-to-island dispersal paradigm. Moreover, our study demonstrates that island systems can act as biodiversity sources—not merely sinks—and provides a rare empirical example of back dispersal in primary freshwater species.

## Introduction

Understanding geographical distribution patterns of species and the processes underlying their formation provides important insights into the mechanism of biodiversity generation (Avise, 2000). In particular, repeated dispersal and subsequent vicariance events play key roles in shaping present-day distribution patterns (Watanabe et al., 2017). During the Pleistocene, sea levels fluctuated repeatedly in response to glacial-interglacial cycles on timescales of tens to hundreds of thousands of years (Avise, 2000; Grant et al., 2014; Spratt & Lisiecki, 2016). The sea level during the Last Glacial Maximum (LGM) was approximately − 125 m from the present (Yokoyama et al., 2018). These sea-level changes repeatedly led to the formation of land bridges during glacial periods and their subsequent submergence into straits during interglacial periods (Hewitt, 2000). The repeated alternation between dispersal via land bridge formation and vicariance due to strait formation provides an ideal framework for understanding species distributions and the processes driving diversification.

The Japanese Archipelago was formed through rifting from the eastern margin of the Eurasian continent approximately 20–15 Ma and reached its present configuration around 5 Ma (Otofuji et al., 1985). During the Pleistocene, sea-level fluctuations led to the formation of land bridges that repeatedly connected the Japanese Archipelago to the continent. These land bridges facilitated the migration of many species to the archipelago (Tojo et al., 2017). Japanese freshwater fishes, including the species studied in this study, are also thought to have dispersed from the continent via such land bridges during glacial periods (Goto, 1994; Watanabe et al., 2017). Proposed dispersal routes include a western route via the Korean Peninsula, a southwestern route via the Ryukyu Islands, a northern route via Sakhalin Island, and a northeastern route via the Kuril Islands (Aoyagi, 1957; Tojo et al., 2017).

Furthermore, recent studies have reported unusual dispersal patterns from the Japanese Archipelago to the continent, termed “back dispersal” (also referred to as reverse colonization) (e.g., Suzuki et al., 2021; Ikeda & Motokawa, 2021). On continental islands formed by the rifting of landmasses from the continent, biota typically originates from a limited subset of continental organisms that become isolated during this process (Ali, 2018). As a result, species diversity on such islands is generally lower than that on the continent (Kier et al., 2009). Nevertheless, endemic species arising through unique evolutionary processes on isolated islands are well documented (Ota, 1998; Tojo et al., 2017). However, given that ecological niches on the continent are already largely occupied, back dispersal from remote islands to the continent has traditionally been regarded as unlikely, and therefore such processes have received comparatively limited attention in biogeographic studies. Understanding these processes is important for re-evaluating the role of island systems as potential sources, rather than sinks, of biodiversity (Bellemain & Ricklefs, 2008; Hutsemékers et al., 2011).

Among these proposed migration routes, including those involving back dispersal, this study focuses on the northern route in the Japanese Archipelago. The Soya Strait, located between Hokkaido and Sakhalin (Fig. 1b), has a reported depth of 50–60 m (Ono & Igarashi, 1991). During the Pleistocene, cyclical changes in sea level led to repeated formation and submergence of land bridges across this strait (Fujimaki, 1994; Goto, 1994). Thus, this region provides an ideal system for investigating the roles of dispersal and vicariance in shaping biodiversity. When the land bridge of the Soya Strait was formed during glacial periods, several primary freshwater fishes, including the pond minnow *Rhynchocypris percnurus sachalinensis*, the Japanese stone loach *Barbatula oreas*, and the eight-barbel loach *Lefua nikkonis*, were suggested to have dispersed from the continent to Hokkaido via Sakhalin (Goto, 1994; Sakai et al., 2014; Ooyagi et al., 2018).

**Fig. 1.**
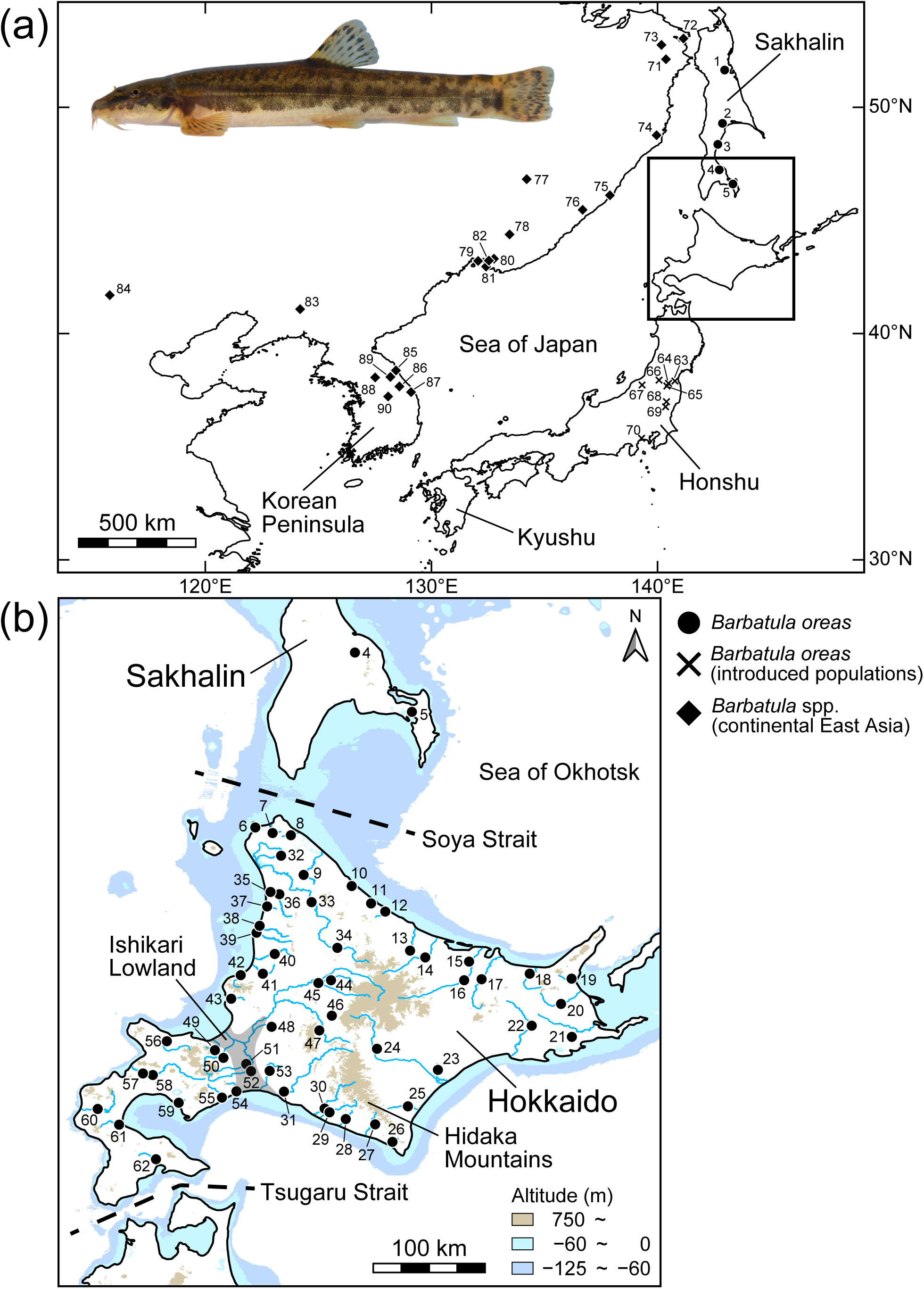
Sampling locations of the genus *Barbatula* species.

In Hokkaido, the uplift of the Hidaka Mountains and the formation of volcanoes occurred resulting from the collision of tectonic plates. The Hidaka Mountains began to uplift 10–5 million years ago (Ono, 1994; Koaze et al., 2003), with the most active uplift occurring during the Quaternary (Kimura & Kusunoki, 1997; Honza, 1999). Moreover, volcanic activity since the Quaternary has led to the distribution of Quaternary volcanoes along the Kuril Arc extending across eastern Hokkaido (Nakano et al., 2013). These processes have resulted in a complex geological history in Hokkaido, involving repeated connections to and separations from the continent, as well as mountain uplift. This history may have promoted population fragmentation and genetic differentiation of freshwater species in Hokkaido. The Japanese stone loach *Barbatula oreas*, the focal species of this study, is the primary freshwater fish distributed from Sakhalin to Hokkaido (Kottelat, 2012; Nakajima & Uchiyama, 2017), making it a suitable model species for discussing historical biogeography and the formation of population genetic structure (Watanabe et al., 2006). This species is widely distributed throughout between upstream and downstream river sections and mainly inhabits lotic environments with substrates ranging from sand and gravels to cobbles and/or boulders (Nakajima & Uchiyama, 2017).

Three primary dispersal processes have been proposed for primary freshwater fishes. First, dispersal through paleo-catchment systems formed during periods of sea-level regression (Hughes et al., 2009; Watanabe & Takahashi, 2010). During such periods, adjacent river systems became connected at their estuaries, forming larger catchment networks. This process is particularly likely in regions with bay-shaped coastal topography. In Japan, genetic exchange via paleo-catchment systems has been reported in areas such as Ise Bay and the Seto Inland Sea (e.g., Watanabe & Nishida, 2003; Watanabe & Mori, 2008; Suzuki et al., 2016; Ito et al., 2019; Ito et al., 2020a; Cho & Mukai, 2023). In contrast, the relationship between paleo-catchment configurations across Hokkaido and Sakhalin and patterns of genetic variation in freshwater species has not been explicitly examined. Second, dispersal between adjacent river systems, which can occur either episodically (e.g., during flooding events) or persistently in low-gradient regions (Watanabe & Takahashi, 2010). In areas with extensive continental shelves, sea-level regression expands downstream floodplains, increasing the likelihood of connectivity among river systems. Such conditions are present around Hokkaido and Sakhalin, where genetic exchange may have occurred (Fig. 1b). Third, dispersal across watersheds driven by geomorphological events such as river capture in headwater regions (Hughes et al., 2009; Watanabe & Takahashi, 2010). *Barbatula oreas*, which is widely distributed throughout river systems, is expected to have expanded its range through these dispersal processes. However, the relative importance of each process in shaping its current distribution remains unclear.

In this study, we investigated the population genetic structure of *B. oreas* using not only mitochondrial DNA (mtDNA) and nuclear DNA (nDNA), but also genome-wide single nucleotide polymorphism (SNP) data. Based on these data, we aimed to: (1) infer the dispersal process across the Soya Strait and its role in diversification, (2) evaluate the impact of geological events on population genetic structure within Hokkaido, and (3) identify the primary dispersal processes contributing to range expansion. In particular, we assessed dispersal direction by integrating ancestral area estimation with patterns of genetic variation between Hokkaido and Sakhalin.

## Materials and Methods

### 1. Sampling

We collected 301 stone loach specimens of the genus *Barbatula* from 90 localities across Hokkaido and Honshu (Japan), Primorsky Krai, Khabarovsky Krai and Sakhalin (Russia), northeastern China, and Korean Peninsula (South Korea) (Fig. 1; Table S1; Table S2). The specimens of mainland East Asia could not be reliably identified at the species level due to insufficient previous taxonomic investigations within the genus *Barbatula*. Therefore, the 42 specimens from Russia, China, and South Korea were treated as *Barbatula* spp. A total of 259 specimens of *Barbatula oreas* were collected from 70 sites on Sakhalin, Hokkaido, and Honshu Islands (Fig. 1; Table S1). Among these, the Honshu populations’ 21 specimens are all considered to have been artificially introduced (Tojo & Hosoya, 1998; Yashima et al., 2011). Almost all specimens were preserved in 99.5% ethanol in the field. Other stone loach species from the families Nemacheilidae and Cobitidae as outgroups were used (Table S3).

Collecting of the target fish species was carried out using a D-frame net.

After excising a small, regenerative portion of the tail fin tissue (a few millimeters square) for genetic analysis, the captured fish was released back into the wild.

In addition, the corresponding author (KT) and GU have completed training on the ethics of animal experimentation at Shinshu University (CertificationNo.2025-0056, 0065), where they are affiliated, and the series of research activities were conducted in accordance with those regulations.

### 2. DNA extraction, amplification, sequencing, and alignment

Total genomic DNA was extracted from ethanol-preserved pectoral fin tissue using a DNeasy Blood & Tissue Kit (Qiagen, Hilden, Germany), following the manufacturer’s protocol. Fragments of the mtDNA, including cytochrome *b* (*cytb*), cytochrome c oxidase subunit I (*COI*), the displacement loop (D-loop), and the 12S rRNA–tRNA-Val–16S rRNA region, and the nDNA, including recombination activating 1 (*RAG1*), were amplified by polymerase chain reaction (PCR). Primers and PCR conditions are provided in Table S4. Ex Taq (Takara, Shiga, Japan) was used for *cytb*, whereas rTaq (TOYOBO, Osaka, Japan) was used for the other regions as the DNA polymerase. PCR products were purified using ExoSAP-IT (GE Healthcare, Amersham, UK). Purified DNA fragments were sequenced using the BigDye Terminator v1.1 Cycle Sequencing Kit (Applied Biosystems, Foster City, CA, USA). Sequencing was performed on an automated DNA sequencer (ABI 3130 or 3130xl DNA Analyzer; Perkin Elmer/Applied Biosystems). All sequences were trimmed and assembled using CLC Workbench software (CLC bio, Aarhus, Denmark) and aligned using MAFFT v.7.222 (Katoh and Standley, 2013). All sequencing data were deposited to the DNA Data Bank of Japan (DDBJ database), and the accession numbers are shown in Table S1 and Table S2.

### 3. Phylogenetic and population genetic analyses based on a sequence-based dataset

The number of haplotypes (Nh), haplotype diversity (Hd), and nucleotide diversity (π) for each clade and subclade were calculated for each mtDNA region (*cytb*, *COI*, D-loop, and 12S rRNA) using DnaSP v.6.12.03 (Rozas et al., 2017). Haplotype networks were constructed based on the mtDNA *cytb* region using the TCS method (Clement et al., 2002) in PopART v.1.7 (Leigh and Bryant, 2015). Pairwise *F*_ST_ values between clades and subclades were calculated using Arlequin v.3.1 (Excoffier et al., 2005). Additionally, analysis of molecular variance (AMOVA) was performed based on the mtDNA dataset using Arlequin. Populations were grouped based on clades as determined by mtDNA and further subdivided into Sakhalin and Hokkaido populations. *P*-values were calculated using nonparametric permutation tests with 10,000 permutations.

Maximum-likelihood (ML) analyses were conducted using IQ-TREE v.2.2.2.7 (Minh et al., 2020) based on each mtDNA and nDNA sequence. The appropriate substitution models for the mtDNA regions were selected using Kakusan4 v.4.0 (Tanabe, 2011) based on the Bayesian Information Criterion (BIC). For the mtDNA dataset, the best-fit models were as follows: K80+G, HKY, and TN93+G for the first, second, and third codon positions of the mtDNA *cytb* region; TN93+G, TN93+I, and F81+I for the first, second, and third codon positions of the mtDNA *COI* region; TN93+G for the mtDNA D-loop region; TVM+G for the mtDNA 12S rRNA region; K80 for the mtDNA tRNA-Val region; and HKY+G for the mtDNA 16S rRNA region. For the nDNA *RAG1* dataset, the best-fit substitution model was K80+G. Node support was assessed using the SH-like approximate likelihood ratio test (SH-aLRT; Guindon et al., 2010) and ultrafast bootstrap (UFBoot; Minh et al., 2013) with 1,000 replicates. SH-aLRT ≥ 80% and UFBoot ≥ 95% are generally considered strong support levels (Guindon et al., 2010; Minh et al., 2013).

Bayesian uncorrelated lognormal relaxed clock analyses based on the mtDNA *cytb* and 12S rRNA regions were performed with BEAST 2 v.2.4.7 (Bouckaert et al., 2014) to estimate divergence time among clades and subclades of *B. oreas*. Due to the lack of fossil evidence for Nemacheilidae, divergence times were estimated using (1) molecular clocks and (2) fossil calibration points. The substitution rates estimated for the genus *Cobitis* (Cobitidae) were used as follows: 0.68% per Myr for the mtDNA *cytb* (Doadrio and Perdices, 2005) and 0.315% per Myr for the mtDNA 12S rRNA (Ludwig et al., 2001). Two fossil records were used: (a) *Cobitis longipectoralis* (18 Ma; Chen et al., 2010) and (b) *Triplophysa opinata* (mid-upper Miocene; Prokofiev, 2007). We treated *Cobitis longipectoralis* as a stem group of *Cobitis sensu lato* following previous studies (Šlechtová et al., 2008; Kim et al., 2013; Perdices et al., 2016; Okada et al., 2024). We used these two calibration points with the log-normal distributions and birth-death tree priors (a: Mean = 1.5, Stdev = 1.66, Offset = 17.5; and b: Mean = 1.5, Stdev = 1.67, Offset = 5.3). The positions of these calibration points are shown in Fig. S1.

The Bayesian analysis and divergence time estimation were performed based on the mtDNA *cytb* and 12S rRNA regions using BEAST 2. TN93+G for the mtDNA *cytb* region and HKY+G for the mtDNA 12S rRNA region were selected based on BIC in Kakusan4. MCMC simulations were run for 200 million generations, sampling every 4,000 generations, with the first 10% discarded as burn-in. The output files were checked by examining the effective sample size (ESS > 200) using Tracer v.1.7.2 (Rambaut et al., 2018). The results were summarized by TreeAnnotator in the BEAST package and visualized using FigTree v.1.4.3 (Rambaut, 2016).

*Barbatula oreas* distributed in Hokkaido and Sakhalin has been treated as a synonym of *Barbatula toni* distributed in the Amur River of Russia (Dyldin & Orlov, 2016). To evaluate species identification, pairwise distance (*p*-distance) between species of the genus *Barbatula* were calculated using MEGA v.10.2.6 (Kumar et al., 2018) based on the mtDNA *cytb* region.

### 4. Demographic analyses

To detect past demographic changes and deviations from neutrality for each clade and subclade of *B. oreas*, based on the mtDNA *cytb* dataset, we conducted mismatch distribution analysis under the sudden expansion model (Rogers and Harpending, 1992), as well as Tajima’s *D* (Tajima, 1989) and Fu’s *Fs* (Fu, 1997) tests using Arlequin.

We also performed Extended Bayesian Skyline Plot (EBSP; Heled and Drummond, 2008) analysis to estimate the demographic history of each subclade using BEAST 2. EBSP analyses were conducted using the mtDNA *cytb* region, 12S rRNA region, and a combined dataset of both regions. The substitution model for each region was selected using Kakusan4. MCMC analyses were run using one billion generations, sampled every 1,000 generations with the first 10% discarded as burn-in. A strict molecular clock was assumed, with substitution rates of 0.68% for the mtDNA *cytb* region and 0.315% for the mtDNA 12S rRNA region. The output data was checked by inspecting ESS values (ESS > 200) using Tracer v.1.7.2. EBSP log files were plotted using R (R Core Team, 2023).

### 5. Ancestral area reconstruction

To estimate the ancestral range of *B. oreas*, we conducted the Bayesian Binary MCMC (BBM) and Statistical Dispersal–Extinction–Cladogenesis (S-DEC) methods using RASP v.4.4 (Yu et al., 2020). BEAST-generated trees were used as input files for these analyses. Six different areas were assigned based on the species’ distribution: Hokkaido, Sakhalin, the Korean Peninsula, continental East Asia (Russia and China), Kyushu and Honshu, and Europe. BBM analysis was run using 500 million generations, sampled every 1,000 generations.

### 6. GRAS-Di sequencing and SNP genotyping

To obtain genome-wide nuclear SNPs, 133 specimens of the genus *Barbatula* from 74 localities were analyzed using genotyping-by-sequencing (GBS) based on the genotyping by random amplicon sequencing-direct (GRAS-Di method; Hosoya et al., 2019). This method employs a two-step PCR process with random primers for library construction. Library preparation and sequencing were outsourced to GeneBay Co., Ltd. For detailed methodology, refer to the following studies: Hosoya et al. (2019) and Ito et al. (2020b). Briefly, DNA samples were directly amplified using 64 random primers (Table S5), in which individuals were tagged with independent adapter sequences. All amplicons were sequenced using MGISEQ-G2000RS (DNBSEQ-G400RS). The raw sequencing data have been deposited in DDBJ (Table S1; Table S2).

Quality checking and primer trimming of the raw reads were performed using fastp v.0.23.4 (Chen et al., 2018) while removing bases with a quality score below 20 and reads shorter than 100 bp (-3 -q 20 -l 100). *De novo* assembly and SNP calling were conducted using the *denovo_map.pl* program in Stacks v.2.66 (Catchen et al., 2011) with parameters (-M 3 -m 3). SNP filtering was performed using the *populations* program in Stacks.

Furthermore, the reads processed by fastp were mapped to the reference genome of *Barbatula barbatula* (GCA_037178815.1; Laczkó et al., 2025) using BWA-MEM2 v.2.2.1 (Vasimuddin et al., 2019) with the default parameters. The output SAM files were sorted and converted to BAM format using SAMtools v.1.21 (Li et al., 2009). Variant calling and joint genotyping were conducted using the *HaplotypeCaller* and *GenotypeGVCFs* programs in Genome Analysis Toolkit (GATK) v.4.6.1 (McKenna et al., 2010) with parameters (-ERC GVCF, -stand-call-conf 20). The resulting variant datasets were filtered using VCFtools v.0.1.16 (Danecek et al., 2011). Detailed filtering criteria for each analysis are provided in Table S5.

### 7. Phylogenetic and population genetic analyses based on nuclear SNP dataset

Genome-wide heterozygosity of each individual was estimated using ANGSD v.0.940 (Korneliussen et al., 2014) based on the BAM files after removing duplicate reads with SAMtools, with quality filtering (-minQ 30, - minMapQ 10) and depth filtering (-setMinDepth 8, -setMaxDepth 300).

Population structure based on nuclear SNPs data was inferred using ADMIXTURE v.1.3.0 (Alexander et al., 2009). The resulting Q matrices were evaluated using evalAdmix v.0.962 (Garcia-Erill & Albrechtsen, 2020). In addition, we performed clustering using the Discriminant Analysis of Principal Components (DAPC) program of the R package adegenet v.2.1.1 (Jombart, 2008). Phylogenetic networks were constructed using the Neighbor-Net algorithm in SplitsTree4 v.4.19.1 (Huson and Bryant, 2006) with Kimura’s 2-parameter (K2P) distances (Kimura, 1980). PCA was performed using Tassel v.5.2.90 (Bradbury et al., 2007). Pairwise *F*_ST_ values between clusters were calculated using Arlequin. Additionally, analysis of molecular variance (AMOVA) was performed based on the nuclear SNPs dataset using Arlequin. Populations were grouped based on clusters as determined by nuclear SNP and further subdivided into Sakhalin and Hokkaido populations.

Maximum-likelihood (ML) analyses were conducted using IQ-TREE based on concatenated SNP alignment generated from the nuclear SNPs dataset. The model selection for the concatenated SNP alignment was performed using Ascertainment bias correction (ASC) model (-m MFP+ASC).

To infer phylogenetic relationships among clusters, we conducted phylogenetic analyses using SVDquartets method (Chifman & Kubatko, 2014) of PAUP* v.4.0a169 (Swofford, 1998). Lineage D–F were designated as outgroup, 500,000 quartets were randomly sampled, node support was evaluated by 1,000 bootstrap replicates.

### 8. Gene flow analyses

To estimate recent gene flow patterns among clusters, directional relative migration rates were inferred using the divMigrate function of the R package diveRsity v.1.9.90 (Keenan et al., 2013) based on Jost’s *D* with 1,000 bootstrap replicates.

### 9. Visualization of paleo-drainage systems

To visualize paleo-drainage systems around Sakhalin and Hokkaido during glacial periods, channel network analysis was performed using SAGA GIS v.7.8.2 (Conrad et al., 2015). The GEBCO_2023 grid (GEBCO Compilation Group, 2023), a 15-arc-second resolution dataset covering both land and ocean, was used as input. The tool Fill Sinks XXL (Wang and Liu, 2006) was used to identify and fill surface depressions in the digital elevation models. The paleo-drainage networks were generated by setting the threshold to 4 and 5 using the tool Channel Network and Drainage Basins.

## Results

### 1. Phylogenetic analysis and species identification based on mtDNA and nDNA

We obtained fragments of the following lengths: 553 bp of the mtDNA *cytb* region; 575 bp of the mtDNA *COI* region; 910 bp of the mtDNA D-loop region; 1,214 bp of the mtDNA 12S rRNA, tRNA-Val, and 16S rRNA regions; 862 bp of the nDNA *RAG1* region. Based on the mtDNA *cytb* region of *B. oreas* from Sakhalin and Hokkaido Islands, a total of 97 haplotypes were detected, having high haplotype diversity (Hd = 0.964) and nucleotide diversity (π = 0.025; Table 1). In Honshu populations, which are considered to have been artificially introduced, two haplotypes (s34, s50) shared with Hokkaido populations were detected (Fig. 2; Table S1). Populations in Sakhalin Island and west of the Ishikari Lowland possessed unique haplotypes not shared with Hokkaido populations (Fig. 2; Table S1). Among regions, genetic diversity was highest in the mtDNA D-loop region and lowest in the mtDNA 12S rRNA region (Table S6). Phylogenetic analysis indicated that *B. oreas* forms two major clades (Northern and Southern Clades) including six subclades (N1–N2, S1–S4; Fig. 2). Additionally, pairwise *F*_ST_ values between clades and subclades were approximately 0.7 or higher, indicating that clades and subclades were highly differentiated (Table S7). Genetic diversity within the clades and subclades was also high (Hd = 0.758–0.939, π = 0.002–0.014; Table 1).

**Fig. 2.**
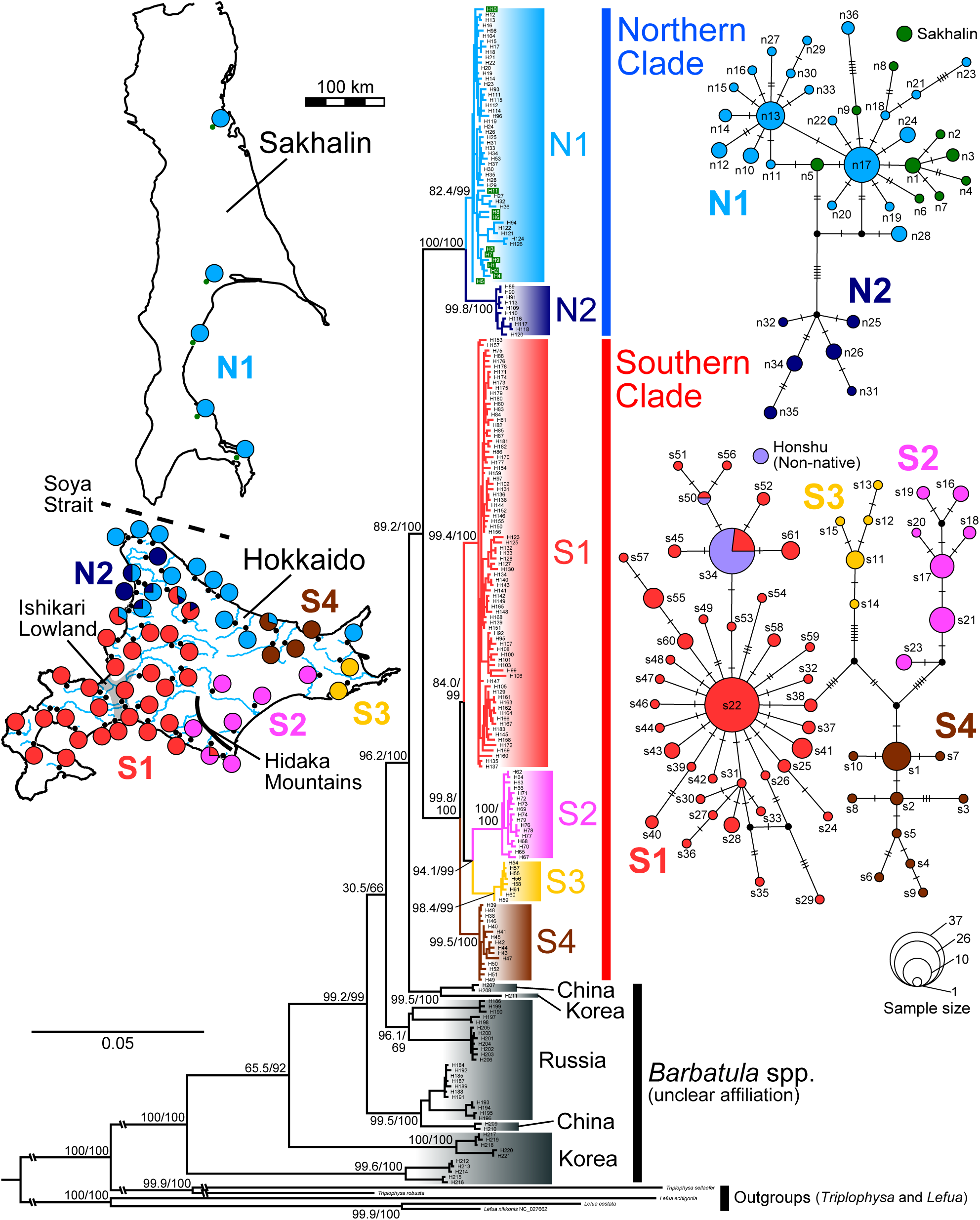
Left: Maximum likelihood tree of the genus *Barbatula* and the related species based on the mtDNA sequence 3,252 bp (*cytb*, 553 bp; *COI*, 575 bp; D-loop, 910 bp; 12S rRNA, tRNA-Val and 16S rRNA, 1,214 bp). The numbers at major nodes indicate SH-aLRT (left) and UFBoot (right) values. Green indicates the haplotypes detected in Sakhalin. A map shows the distribution of each subclade in *Barbatula oreas*. Right: Haplotype networks of *Barbatula oreas* based on the mtDNA *cytb*. Circle size reflects the number of specimens sharing each haplotype, and small black circles indicate missing haplotypes. The haplotypes detected in the Sakhalin and Honshu populations are shown as pie charts in green and purple, respectively.

**Table 1.**
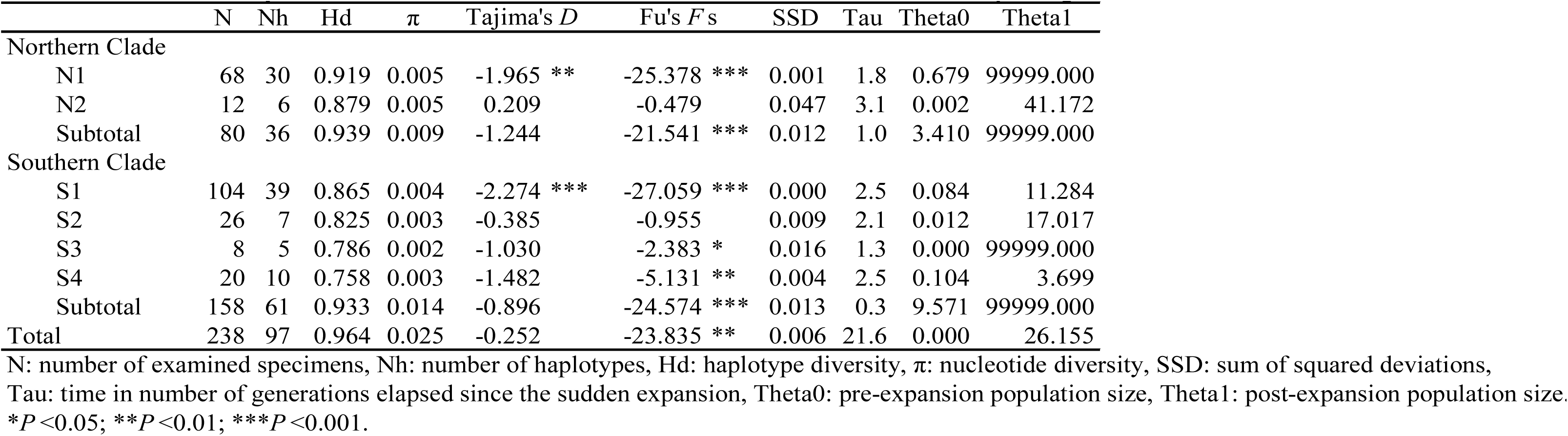
Genetic diversity of *Barbatula oreas* in Hokkaido and Sakhalin based on the mtDNA *cytb* region.

| | N | Nh | Hd | $\pi$ | Tajima's *D* | Fu's *F*<sub>s</sub> | SSD | Tau | Theta0 | Theta1 |
| --- | --- | --- | --- | --- | --- | --- | --- | --- | --- | --- |
| Northern Clade |  |  |  |  |  |  |  |  |  |  |
| N1 | 68 | 30 | 0.919 | 0.005 | -1.965 ** | -25.378 *** | 0.001 | 1.8 | 0.679 | 99999.000 |
| N2 | 12 | 6 | 0.879 | 0.005 | 0.209 | -0.479 | 0.047 | 3.1 | 0.002 | 41.172 |
| Subtotal | 80 | 36 | 0.939 | 0.009 | -1.244 | -21.541 *** | 0.012 | 1.0 | 3.410 | 99999.000 |
| Southern Clade |  |  |  |  |  |  |  |  |  |  |
| S1 | 104 | 39 | 0.865 | 0.004 | -2.274 *** | -27.059 *** | 0.000 | 2.5 | 0.084 | 11.284 |
| S2 | 26 | 7 | 0.825 | 0.003 | -0.385 | -0.955 | 0.009 | 2.1 | 0.012 | 17.017 |
| S3 | 8 | 5 | 0.786 | 0.002 | -1.030 | -2.383 * | 0.016 | 1.3 | 0.000 | 99999.000 |
| S4 | 20 | 10 | 0.758 | 0.003 | -1.482 | -5.131 ** | 0.004 | 2.5 | 0.104 | 3.699 |
| Subtotal | 158 | 61 | 0.933 | 0.014 | -0.896 | -24.574 *** | 0.013 | 0.3 | 9.571 | 99999.000 |
| Total | 238 | 97 | 0.964 | 0.025 | -0.252 | -23.835 ** | 0.006 | 21.6 | 0.000 | 26.155 |

N: number of examined specimens, Nh: number of haplotypes, Hd: haplotype diversity, $\pi$ : nucleotide diversity, SSD: sum of squared deviations, Tau: time in number of generations elapsed since the sudden expansion, Theta0: pre-expansion population size, Theta1: post-expansion population size. \* $P < 0.05$ ; \*\* $P < 0.01$ ; \*\*\* $P < 0.001$ .

The ML tree based on the six mtDNA regions analyzed indicated that *B. oreas* populations form a monophyletic group (Fig. 2), divided into two major clades: the Northern Clade (Sakhalin and Northern Hokkaido populations) and Southern Clade (Southern Hokkaido populations). The broadly similar northern-southern genetic structure was also recovered from the ML tree based on the nDNA *RAG1* region (Fig. S2), although several mito-nuclear discordances were observed. Several localities contained specimens from both major clades. The Subclades N1, S1, and S2 were distributed across major geographic barriers such as the Soya Strait, Ishikari Lowland, and Hidaka Mountains, respectively. Continental East Asian populations of *Barbatula* spp. from Russia, China, and South Korea form a paraphyletic group to *B. oreas* populations.

In addition, genetic distances (*p*-distance) based on the mtDNA *cytb* region between *B. oreas* populations and continental East Asian species were roughly comparable to distances between recognized *Barbatula* species (Table S8).

### 2. Divergence time estimation and ancestral range reconstruction based on mtDNA

We inferred the divergence times using two approaches: (1) molecular clocks and (2) fossil calibration points. Both methods estimated broadly similar divergence times (Table S9). For downstream analyses, we adopted the fossil calibration method.

In the time-calibrated tree, the divergence time between *B. oreas* and its congeneric species from China and Korea was estimated to be 3.20 Ma [95% highest posterior density (HPD): 4.55–2.00 Ma] (Fig. 3). The divergence time between the Northern and Southern Clades was estimated to be 2.32 Ma (95% HPD: 3.33–1.40 Ma) (Fig. 3; Table S9). The divergence time between Subclades N1 and N2 was estimated to be 1.04 Ma (95% HPD: 1.60–0.57 Ma), and the divergence time between Subclades S1+S2+S3 and S4 was 1.41 Ma (HPD: 2.04–0.82 Ma).

**Fig. 3.**
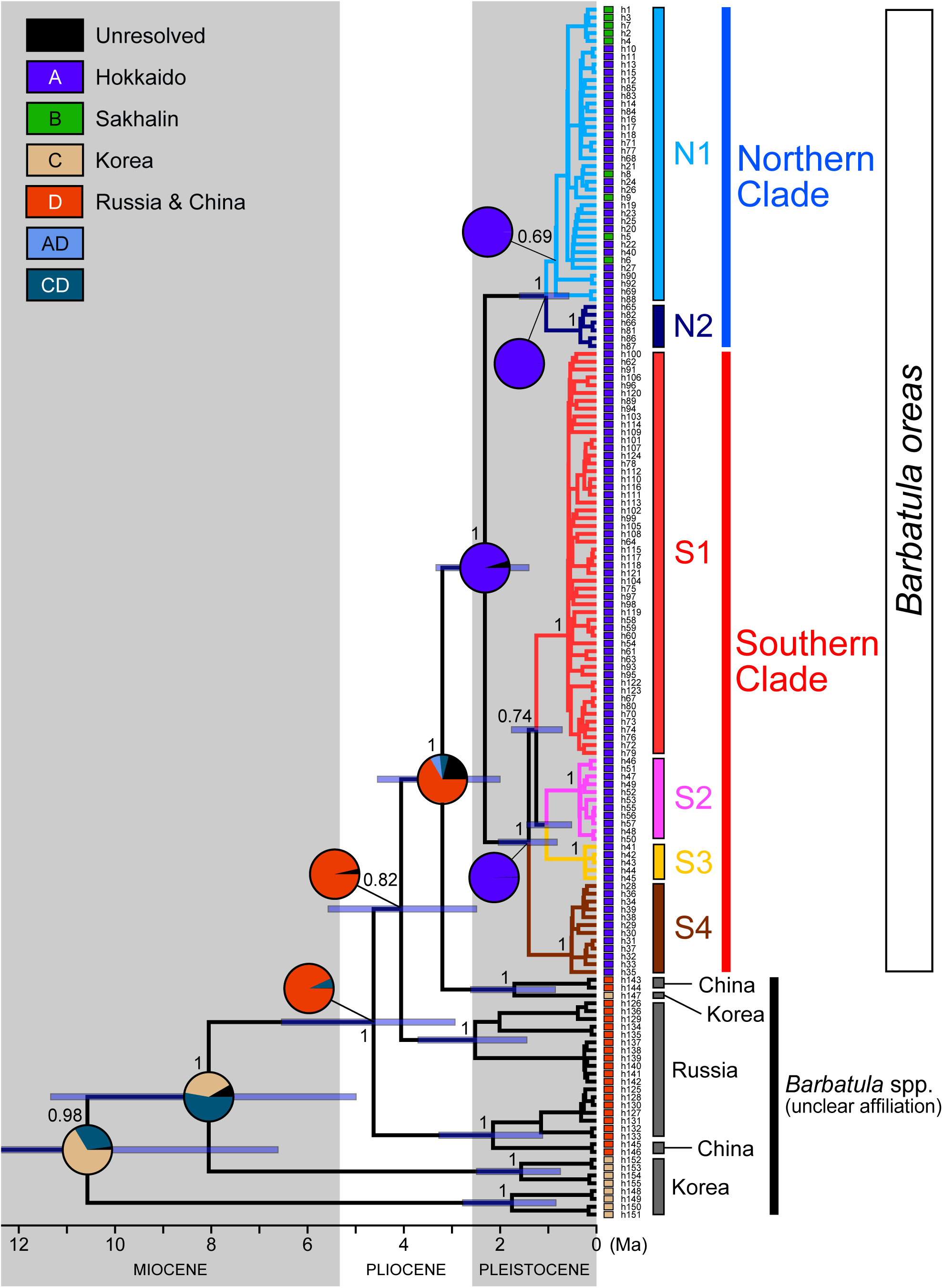
Time-calibrated Bayesian tree of the genus *Barbatula* inferred using the mtDNA *cytb* and 12S rRNA sequences, with calibration points derived from the fossil record. The numbers near major nodes indicate the Bayesian posterior probabilities, and the node bar indicates 95% highest density confidence intervals of divergence times. Ancestral area reconstruction was performed using Bayesian Binary MCMC (BBM). The pie charts indicate probabilities of ancestral areas.

Ancestral ranges of *Barbatula* species were inferred using BBM and S-DEC (Fig. 3; Fig. S3). Both analyses indicated that the ancestral area of *B. oreas* was Hokkaido. Similarly, the ancestral area of the Northern Clade and Subclade N1 was inferred to be Hokkaido, suggesting dispersal of *B. oreas* from Hokkaido to Sakhalin. Furthermore, the results suggested that the ancestral species of the genus *Barbatula* originated in continental East Asia.

### 3. Population structure and demographic history based on mtDNA

Haplotype networks were constructed based on the mtDNA *cytb* region for each major clade (Fig. 2) and were divided into subclades like for phylogenetic analyses. Star-shaped networks were observed in Subclades N1 and S1. Haplotypes in the Sakhalin populations were unique; interestingly, these haplotypes diverged multiple times from those of Hokkaido populations.

To detect past demographic expansions, neutrality tests and mismatch distribution analyses were performed based on the mtDNA *cytb* region (Table 1; Fig. 4a). In Subclades N1 and S1, both Tajima’s *D* and Fu’s *Fs* values were significantly negative, and unimodal-shape mismatch distributions were confirmed, indicating recent population expansion. Fluctuation patterns of effective population sizes were further estimated using EBSP analyses based on the mtDNA *cytb* and 12S rRNA regions (Fig. 4b; Fig. S4). These analyses revealed sudden population expansions occurring approximately 0.3–0.1 Ma in Subclades N1 and S1.

**Fig. 4.**
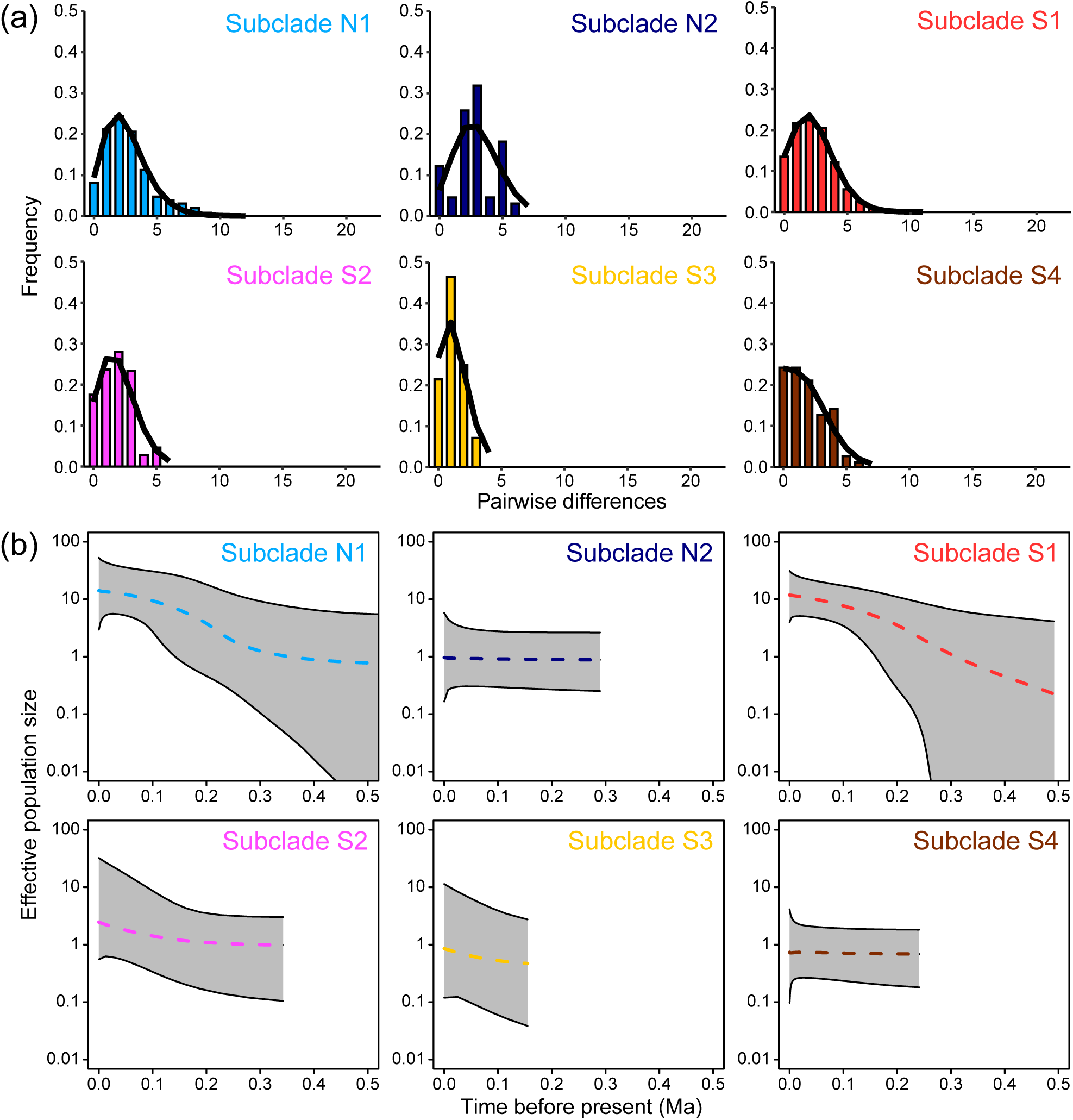
(a) Mismatch distribution analyses for each clade and subclade of *Barbatula oreas* based on the mtDNA *cytb* sequences. Observed distributions of pairwise differences are shown as vertical bars, and expected distributions under the demographic expansion model are shown as black lines. (b) Extended Bayesian Skyline Plots (EBSP) for each subclade based on the mtDNA *cytb* sequences. The dashed lines represent median estimates of the effective population size, and the gray areas indicate 95% highest posterior density intervals.

### 4. Phylogenetic analysis and population structure based on nuclear SNPs

To examine the nuclear genetic structure of the genus *Barbatula*, we conducted ADMIXTURE analyses and constructed the corresponding phylogenetic networks. Results from ADMIXTURE analyses based on both the *de novo* and mapping datasets indicated broadly similar cluster patterns (Fig. S5–S7). Similarly, phylogenetic networks based on both datasets showed no major differences (Fig. 5c–d; Fig. S8). From these results, we selected the mapping dataset for downstream analyses because it provided higher coverage and accuracy.

**Fig. 5.**
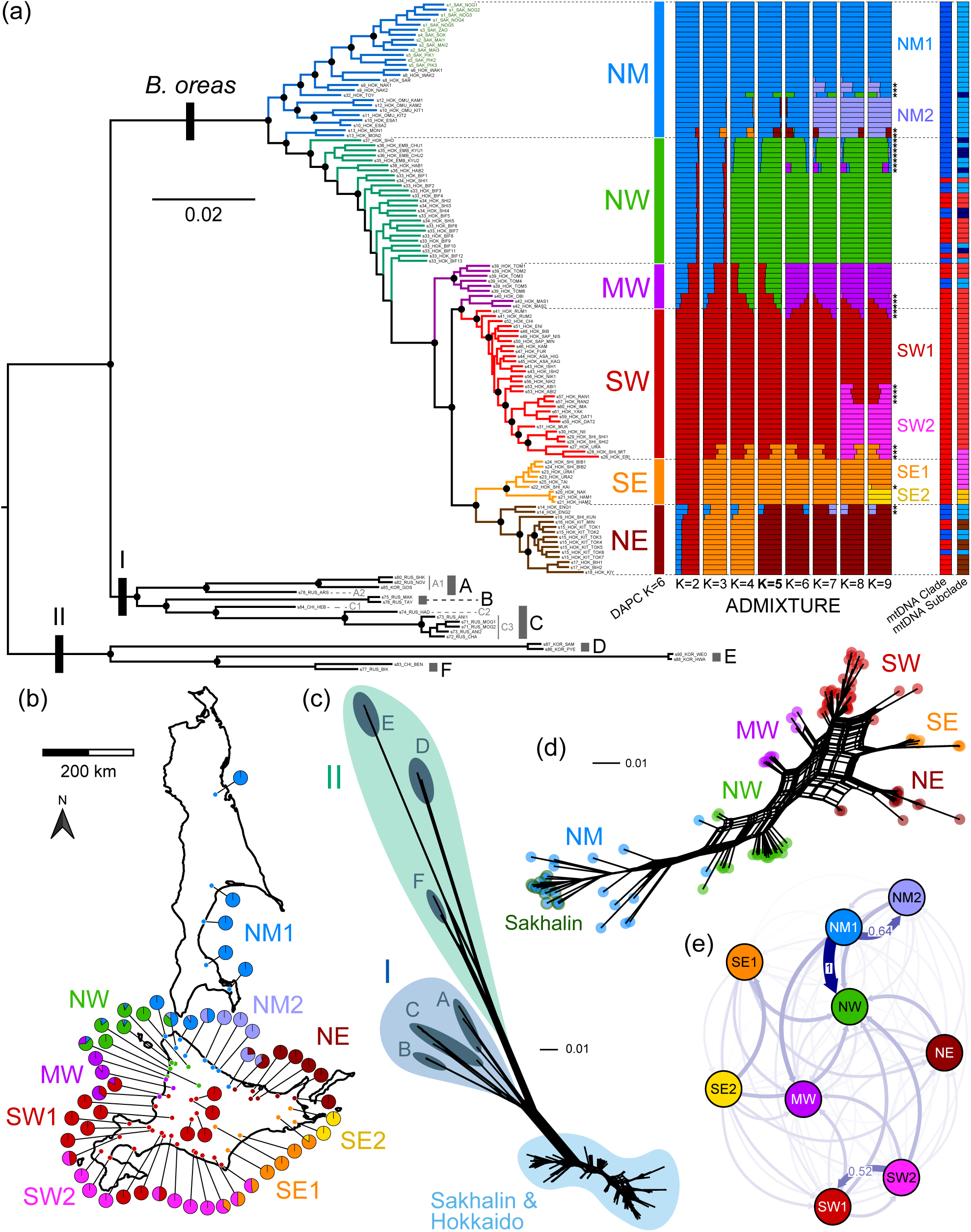
(a) Left: Maximum likelihood tree of the genus *Barbatula* based on 132,537 nuclear SNPs. The black circles indicate nodes with SH-aLRT ≥ 80% and UFBoot ≥ 95%. The specimens from Sakhalin populations are shown in dark green. Right: Population genetic structure of *Barbatula oreas* based on 13,125 nuclear SNPs. From left to right, clustering using DAPC (K = 6), population structure inferred using ADMIXTURE (K = 2–9; the most likely K is shown in bold), and clades and subclades based on mtDNA. Individuals marked with asterisks in ADMIXTURE at K = 9 represent “admixed individuals” among subclusters. (b) Geographic distribution of major clusters and subclusters in *B. oreas*. The colored points indicate major cluster assignment, and pie charts show subcluster composition inferred using ADMIXTURE at K = 9. (c–d) Phylogenetic networks constructed using the Neighbor-net algorithm based on (c) 159,567 SNPs from all samples and (d) 111,035 SNPs from *B. oreas* only. The specimens from Sakhalin populations are shown as dark green circles. (e) Directional gene flow among subclusters estimated using divMigrate based on 8,296 nuclear SNPs. All inferred directional migration links are shown, whereas only migration rates above a threshold of 0.5 are labeled.

After joint genotyping, 5,553,916 sites were obtained from the all-samples dataset (133 samples), and 3,804,291 sites from the *B. oreas* dataset (114 samples). Following filtering, 159,567 SNPs were identified in the all-samples dataset, and 111,035 SNPs in the *B. oreas* dataset. All mapping datasets were covered at average depths ranging from 10x to 300x.

The ML tree constructed from the nuclear SNPs dataset with invariant sites removed indicated that *B. oreas* formed a monophyletic clade, consistent with the mtDNA results, and was genetically distant from continental species lineages (Fig. 5a). Phylogenetic networks also showed a clear separation between *B. oreas* populations and continental lineages (Fig. 5c). Based on these results, we divided the all-samples dataset into *B. oreas* and continental East Asian datasets for subsequent analyses.

To investigate the population genetic structure of *B. oreas*, we conducted clustering analyses based on the *B. oreas* dataset. ADMIXTURE analysis at K = 2 indicated *B. oreas* populations were divided into Northern and Southern lineages (Fig. 5a; Fig. S6). This Northern–Southern division was largely consistent with the Northern and Southern mtDNA clades, although mito-nuclear discordance was observed (Fig. 5a). Cross-validation error values based on ADMIXTURE analyses were relatively low, K = 3–9 (Fig. S9; Table S10). BIC values based on the DAPC method reached a minimum of K = 6 (Fig. S10). Accordingly, the *B. oreas* populations were divided into six clusters: NM (northernmost), NW (northwest), MW (middle west), SW (southwest), SE (southeast), and NE (northeast) (Fig. 5a). These clusters showed distinct geographic distributions, and the NM Cluster was distributed across Sakhalin and Northern Hokkaido (Fig. 5b). In addition, pairwise *F*_ST_ values between the six clusters were approximately 0.1 or higher, suggesting significant genetic differentiation (Table 2a). We thereafter referred to these six clusters as “major clusters”. In ADMIXTURE among the results where K = 6, some individuals possessed genetic components from multiple major clusters (Fig. 5a). In the PCA plot, PC1 separated Northern and Southern populations, while PC2 separated Western and Eastern populations (Fig. S11). Overall, the PCA plot showed a triangular distribution corresponding to Northern, Western and Eastern lineages, where each major cluster occupied a distinct position. Notably, the ML tree based on nuclear SNPs indicated that the Hokkaido populations in the NM Cluster branched at the base of the tree, followed by the Sakhalin populations (Fig. 5a).

**Table 2.** Pairwise *F*_ST_ values (a) between the six major clusters and (b) among the nine subclusters based on 13,125 nuclear SNPs of *Barbatula oreas*.

| (a) |  |  |  |  |  |  | (b) |  |  |  |  |  |  |  |  |  |
| --- | --- | --- | --- | --- | --- | --- | --- | --- | --- | --- | --- | --- | --- | --- | --- | --- |
|  | NM | NW | MW | SW | SE | NE |  | NM1 | NM2 | NW | MW | SW1 | SW2 | SE1 | SE2 | NE |
| NM |  |  |  |  |  |  | NM1 |  |  |  |  |  |  |  |  |  |
| NW | 0.098 |  |  |  |  |  | NM2 | 0.130 |  |  |  |  |  |  |  |  |
| MW | 0.178 | 0.100 |  |  |  |  | NW | 0.146 | 0.122 |  |  |  |  |  |  |  |
| SW | 0.226 | 0.151 | 0.120 |  |  |  | MW | 0.236 | 0.212 | 0.120 |  |  |  |  |  |  |
| SE | 0.301 | 0.235 | 0.268 | 0.239 |  |  | SW1 | 0.274 | 0.261 | 0.155 | 0.169 |  |  |  |  |  |
| NE | 0.221 | 0.176 | 0.210 | 0.209 | 0.251 |  | SW2 | 0.309 | 0.301 | 0.183 | 0.218 | 0.094 |  |  |  |  |
|  |  |  |  |  |  |  | SE1 | 0.361 | 0.361 | 0.240 | 0.322 | 0.275 | 0.335 |  |  |  |
|  |  |  |  |  |  |  | SE2 | 0.414 | 0.419 | 0.291 | 0.386 | 0.353 | 0.415 | 0.338 |  |  |
|  |  |  |  |  |  |  | NE | 0.293 | 0.282 | 0.202 | 0.266 | 0.242 | 0.287 | 0.287 | 0.373 |  |

ADMIXTURE results of K = 3–9, which showed low cross-validation error values, were further evaluated using evalAdmix (Fig. S12). evalAdmix evaluates model fit by estimating correlations of residual differences between individuals (Garcia-Erill & Albrechtsen, 2020). If the model is a good fit for the data, the correlation is close to 0. As a result of the evalAdmix evaluation, the correlation for K = 9 was generally close to 0.

ADMIXTURE result of K = 9 indicated that major clusters were further subdivided into smaller clusters (hereafter referred to as subclusters): the NM Cluster was divided into NM1 and NM2 Subclusters, SW Cluster into SW1 and SW2 Subclusters, and SE Cluster into SE1 and SE2 Subclusters (Fig. 5a). The NM1 Subcluster was comprised of populations from Sakhalin and northernmost Hokkaido. Also, pairwise *F*_ST_ values between the nine subclusters were approximately 0.1 or higher (Table 2b). Where K = 9, specimens possessing genetic components from multiple subclusters were observed between neighboring subclusters (Fig. 5b). Individuals with more than 10% ancestry from minor clusters were classified as admixed individuals and excluded from downstream analyses. Phylogenetic relationships among subclusters inferred using the SVDquartets method were largely consistent with those of ML tree (Fig. S13).

Continental East Asian populations were divided into two major lineages (major lineage I and II) based on ADMIXTURE analyses and phylogenetic networks (Fig. 5c; Fig. S7–S8) and were subsequently differentiated into Lineages A–F based on phylogenetic analysis (Fig. 5a). Mito-nuclear discordance was observed: Lineage F was assigned to major lineage II based on the nuclear SNPs, but to major lineage I based on the mtDNA dataset (Fig. S14). Based on the PCA plot and pairwise *F*_ST_ values, Lineages A and C were further subdivided into Sublineages A1–A2 and C1–C3, respectively (Fig. S15; Table S11).

In all AMOVA results based on both mtDNA and nuclear SNPs datasets, the genetic variation between regions (Hokkaido and Sakhalin) was lower than within regions, and the *F*_CT_ values were low (Table 3).

**Table 3.** Result of analysis of molecular variance (AMOVA) of *Barbatula oreas* between Hokkaido and Sakhalin.

| Source of variation | *d.f.* | Sum of squares | Variance components | Percentage of variation | Fixation indices |
| --- | --- | --- | --- | --- | --- |
| All individuals in North Clade from Hokkaido vs. Sakhalin based on mtDNA 6 regions |  |  |  |  |  |
| Among regions | 1 | 36.4 | 0.756 | 7.7 | $F_{CT} = 0.077$ |
| Among populations within regions | 22 | 415.4 | 4.265 | 43.4 |  |
| Within populations | 56 | 269.4 | 4.811 | 48.9 |  |
| Total | 79 | 721.1 | 9.831 |  |  |
| All individuals in Subclade N1 from Hokkaido vs. Sakhalin based on mtDNA 6 regions |  |  |  |  |  |
| Among regions | 1 | 31.5 | 0.956 | 15.0 | $F_{CT} = 0.150^{**}$ |
| Among populations within regions | 19 | 202.9 | 2.398 | 37.7 |  |
| Within populations | 47 | 141.0 | 2.999 | 47.2 |  |
| Total | 67 | 375.4 | 6.353 |  |  |
| All individuals in NM Cluster from Hokkaido vs. Sakhalin based on 13,125 nuclear SNPs |  |  |  |  |  |
| Among regions | 1 | 2813.3 | 39.177 | 4.2 | $F_{CT} = 0.042^{**}$ |
| Among populations within regions | 11 | 16250.3 | 194.766 | 20.6 |  |
| Within populations | 41 | 29090.4 | 709.521 | 75.2 |  |
| Total | 53 | 48153.9 | 943.463 |  |  |
| All individuals in NM1 Subcluster from Hokkaido vs. Sakhalin based on 13,125 nuclear SNPs |  |  |  |  |  |
| Among regions | 1 | 1418.0 | 47.627 | 6.5 | $F_{CT} = 0.065^{*}$ |
| Among populations within regions | 5 | 4921.9 | 89.871 | 12.3 |  |
| Within populations | 25 | 14838.9 | 593.557 | 81.2 |  |
| Total | 31 | 21178.8 | 731.054 |  |  |

\* $P < 0.05$ ; \*\* $P < 0.01$

### 5. Gene flow and genetic diversity based on nuclear SNPs

Asymmetric gene flow analysis using the divMigrate method with a threshold of 0.5 detected gene flow from NM1 to NM2 and NW, and from SW2 to SW1 (Fig. 5e). The other inferred gene flow values were approximately 0.3 or lower (Table S12).

Based on genome-wide heterozygosity values for each cluster, the values were higher than 0.005 (Fig. S16). Among clusters, the NW Cluster showed the highest heterozygosity, followed by the NM and MW clusters.

### 6. Paleo-drainage reconstruction around Hokkaido and Sakhalin

Paleo-drainage systems around Hokkaido and Sakhalin were reconstructed using channel network analysis to a sea level of − 125 m, corresponding to the Last Glacial Maximum (Fig. 6). The reconstructed paleo-drainage networks extended across both Hokkaido and Sakhalin (Fig. 6b). Additionally, two adjacent paleo-drainage systems were identified: one included the Teshio river system (Teshio paleo-drainage) and the other included the Kotanbetsu river system (Kotanbetsu paleo-drainage) (Fig. 6c).

**Fig. 6.**
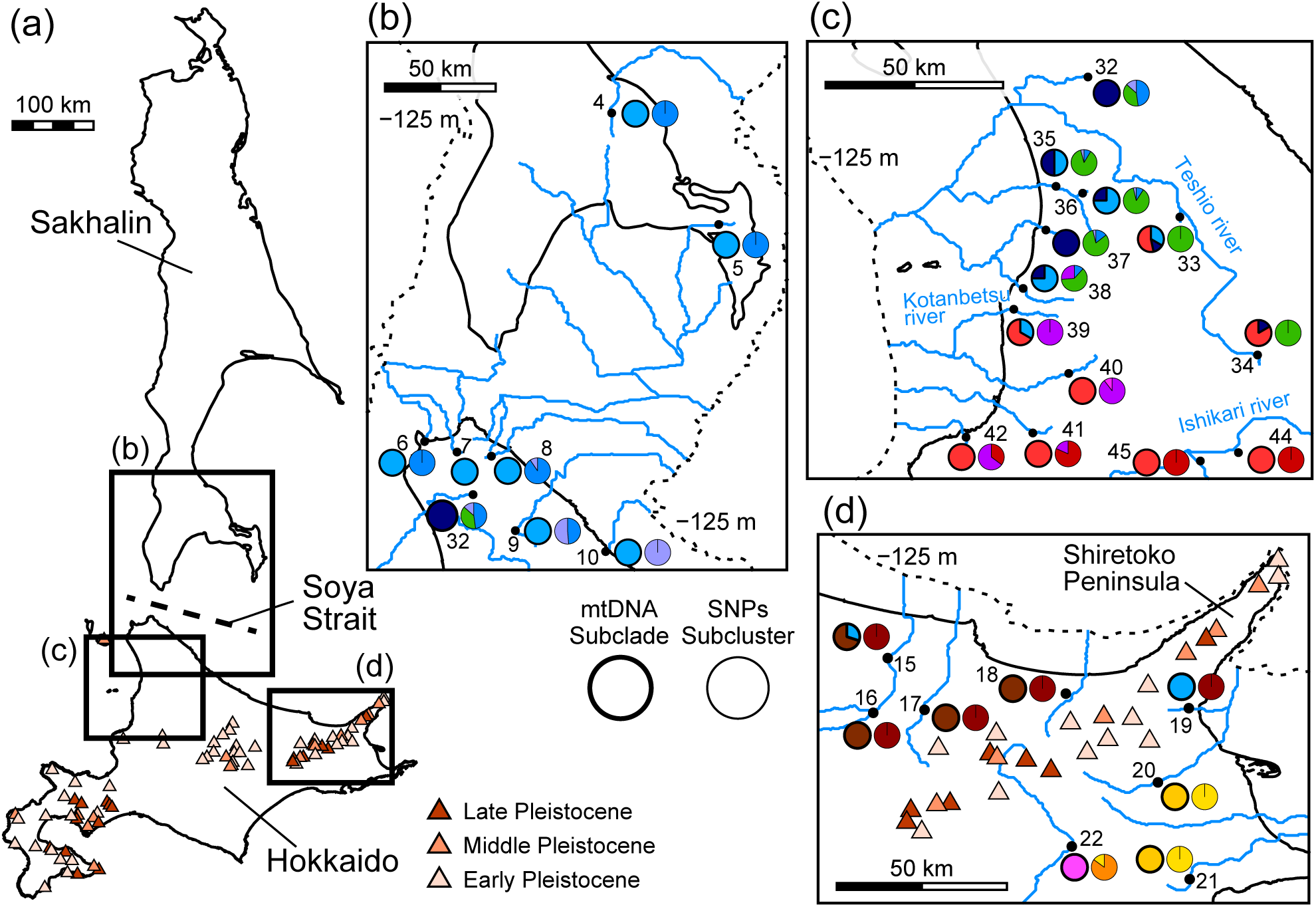
(a) Distribution of Quaternary volcanoes in Hokkaido and Sakhalin. A map shows the volcanoes in different colors according to the timing of their initial activity. (b–d) Regional maps showing reconstructed paleo-drainage systems inferred using SAGA GIS. Thick-lined pie charts (left) indicate the distribution of each subclade based on mtDNA, whereas thin-lined pie charts (right) indicate subcluster composition inferred using ADMIXTURE at K = 9. Black dashed lines represent sea level at the last glacial maximum (LGM; − 125 m).

## Discussion

This study revealed the population genetic structure and phylogeographic history of *B. oreas*, providing new insights into dispersal processes in East Asia. *Barbatula oreas* populations were clearly divided into Northern and Southern lineages and were subdivided into regional subclades and subclusters across Hokkaido. In contrast, mito-nuclear discordance and genetic admixture were observed among lineages in Hokkaido. Furthermore, based on ancestral area reconstruction, an unusual northward dispersal process from Hokkaido to Sakhalin was suggested, which contrasts with the commonly assumed southward colonization from the continent via Sakhalin to Hokkaido.

### 1. Species identification and phylogenetic relationships

Phylogenetic analysis and divergence time estimation revealed that *B. oreas* formed a distinct monophyletic group, clearly diverged from continental congeneric species in the late Pliocene (3.20 Ma [95% HPD: 4.55–2.00 Ma]). Ancestral area reconstruction suggested that the genus *Barbatula* originated in continental East Asia, consistent with Šlechtová et al. (2024). Subsequently, *B. oreas* was inferred to have dispersed from continental East Asia to Hokkaido: this dispersal pattern is widely reported in the fauna and flora of Hokkaido (e.g., Fujii et al., 1999; Hirata et al., 2013; Tojo et al., 2017; Ooyagi et al., 2018). Furthermore, although the occurrence of *B. oreas* on Sakhalin Island has been previously reported based on an extremely small sample size (Yatsuyanagi et al., 2024; Tsuji et al., 2026), sampling in this study revealed that *B. oreas* was widely distributed across Sakhalin Island. Including areas with artificial introductions, *B. oreas* extensively ranges across Sakhalin, Hokkaido, and Eastern Honshu (Nakajima & Uchiyama, 2017).

Genetic distance (*p*-distance) values indicated that *Barbatula oreas* was genetically distinct from *Barbatula toni*, with which it has been synonymized (Dyldin & Orlov, 2016). This result supports previous taxonomic treatments (Kottelat, 2012; Nakajima & Uchiyama, 2017) recognizing *Barbatula oreas* as a distinct species.

For continental East Asian species, mito-nuclear discordance was observed, indicating introgression and incomplete lineage sorting among continental species (e.g., Funk & Omland, 2003; Toews & Brelsford, 2012; Yoshida et al., 2026). Thus, for taxonomic clarification of the genus *Barbatula*, an integrative approach incorporating not only mtDNA but also nuclear DNA is needed.

### 2. Geographic genetic structure of *B. oreas*

Phylogenetic and clustering analyses based on both mtDNA and nuclear SNPs consistently identified two genetically differentiated lineages, showing a Northern-Southern geographic separation within Hokkaido. Such a Northern-Southern genetic structure has rarely been reported in other freshwater taxa in Hokkaido; instead, these taxa generally show patterns of Eastern-Western genetic differentiation (e.g., Koizumi et al., 2012; Sakai et al., 2014). One possible geographic factor underlying this Northern-Southern differentiation is the presence of two adjacent paleo-drainage systems (the Teshio-gawa and Kotanbetsu-gawa paleo-catchment) inferred by our paleo-catchment reconstructions, which likely remained separated even during periods of sea-level regression (Fig. 6c). The Northern mtDNA clade and the NW nuclear SNP cluster were primarily distributed in the Teshio-gawa paleo-catchment, whereas the Southern mtDNA clade and the MW nuclear SNP cluster were primarily distributed in the Kotanbetsu-gawa paleo-catchment. This distribution pattern likely reflects connectivity of lower river basins resulting from the form of paleo-catchment systems (e.g., Hughes et al., 2009; Watanabe & Takahashi, 2010). However, some exceptions were observed, particularly near the boundary between the Teshio-gawa and Kotanbetsu-gawa paleo-catchment. In this area, individuals with genetic components of the MW cluster were detected within the Teshio-gawa paleo-catchment and individuals assigned to the Northern clade within the Kotanbetsu-gawa paleo-catchment. In addition, individuals assigned to the Southern clade were detected in the upper reaches of the Teshio-gawa River. The former pattern may be explained by temporary connections between adjacent river systems during events such as floods (e.g., Watanabe & Takahashi, 2010), whereas the latter is likely attributable to river capture from the Ishikari-gawa River into the Teshio-gawa River (e.g., Hughes et al., 2009; Watanabe & Takahashi, 2010; Masuda et al., 2023). Another possible geographic factor underlying this Northern-Southern differentiation is a chain of Quaternary volcanoes (Fig. 6d). The mtDNA of Subclades N1 and S4, and the NE nuclear SNP cluster were primarily distributed north of the chain, whereas the mtDNA of Subclades S2 and S3, and the SE nuclear SNP cluster were primarily distributed south of the chain. However, individuals from site 19 were detected across the barrier. This dispersal may have occurred prior to the formation of the Shiretoko Peninsula, which originated from submarine volcanic eruptions (Goto et al., 2000). Nevertheless, these barriers do not strictly divide the Northern and Southern lineages.

Phylogenetic analysis and divergence time estimation based on mtDNA indicated that *B. oreas* comprises two clades and six subclades that diverged during the Pleistocene. Based on nuclear SNPs, six major clusters and nine subclusters were identified through clustering analyses. As these clades and clusters are geographically separated within Hokkaido, this intraspecific genetic structure likely resulted from allopatric divergence. While the mtDNA-based genetic structure identified in this study is broadly consistent with recent environmental DNA (eDNA) studies (Yatsuyanagi et al., 2024; Tsuji et al., 2026), the use of longer mtDNA sequences and genome-wide SNP datasets in this study provides improved resolution and more reliable inference of phylogeographic patterns.

Furthermore, these clades and clusters exhibit high genetic diversity, which likely reflects the persistence of populations in multiple refugia during the Pleistocene (e.g., Koizumi et al., 2012; Kawai et al., 2013). Such allopatric differentiation is likely associated with refugial isolation during glacial-interglacial cycles (Hewitt, 2000; Stewart et al., 2010).

By contrast, mito-nuclear discordance and mixed ancestry patterns were detected among clades and clusters within Hokkaido, likely reflecting historical secondary contact. In ADMIXTURE results of K = 9, individuals from multiple localities showed genetic components from adjacent subclusters. In contrast, divMigrate analysis detected only limited recent gene flow. These results suggest that river rearrangement during glacial-interglacial cycles may repeatedly have facilitated historical dispersal and population contact among neighboring populations (e.g., Keogh et al., 2025), whereas recent gene flow has been restricted.

### 3. Back dispersal and phylogeographic history of *B. oreas*

Since the ancestral area of the genus *Barbatula* was inferred to be continental East Asia, *B. oreas* has been suggested to have dispersed from continental East Asia to Hokkaido. Given the limited dispersal ability of freshwater fishes and the dispersal scenarios proposed for several northern freshwater taxa (Goto, 1994; Sakai et al., 2014; Ooyagi et al., 2018), this dispersal may have occurred via Sakhalin. However, our ancestral area reconstruction revealed that the Northernmost lineage of *B. oreas* dispersed from Hokkaido to Sakhalin, indicating back dispersal from islands toward the mainland. Although Sakhalin is an island, the Mamiya (Tatar) Strait between Sakhalin and the Eurasian continent is extremely shallow (less than 10 m deep) and may at times allow terrestrial connectivity due to tidal fluctuations or seasonal sea ice formation (Ono & Igarashi, 1991). Therefore, this dispersal process from Hokkaido to Sakhalin can be regarded as island-to-continent back dispersal in a biogeographic context. Such back dispersal is an important but often overlooked process in biogeography, highlighting the potential role of island systems as sources of continental diversity (Bellemain & Ricklefs, 2008; Esposito & Prendini, 2019). Phylogenetic analyses based on both mtDNA and nuclear SNPs consistently indicated that Sakhalin populations are derived from Hokkaido populations, further supporting this dispersal pattern. Given that the divergence time between Subclades N1 and N2 was estimated to be 1.04 Ma (95% HPD: 1.60–0.57 Ma), this back dispersal likely occurred relatively recently. Moreover, EBSP analyses indicated a sudden population expansion in Subclade N1 at approximately 0.3–0.1 Ma, suggesting that demographic expansion may have facilitated the back dispersal event. This result shows a similar temporal and directional pattern to recent botanical studies, which have reported northward dispersal from the Japanese Archipelago to Beringia in the Northern Pacific region (Ikeda et al., 2018, 2020; Kurata et al., 2022).

AMOVA results consistently showed low levels of genetic variation between Hokkaido and Sakhalin, based on both mtDNA and nuclear SNPs, indicating genetic exchanges between these regions. Additionally, our paleo-catchment reconstruction suggested the presence of catchment systems connecting Hokkaido and Sakhalin across the Soya Strait (Fig. 6b). Because *B. oreas* inhabits a wide range of riverine habitats from the upper to lower reaches of rivers (Nakajima & Uchiyama, 2017), the paleo-catchment systems may have offered dispersal corridors enabling *B. oreas* to cross the Soya Strait during glacial periods. Haplotype networks also suggest that Sakhalin haplotypes were derived multiple times from Hokkaido haplotypes, further supporting exchanges between the two regions.

Although our results suggest natural back dispersal from Hokkaido to Sakhalin, recent genetic studies have demonstrated that two loach species in Sakhalin were artificially introduced from Hokkaido (*Lefua nikkonis*, Machida et al., 2021; *Misgurnus* sp. Type I, Okada et al., 2024), raising the possibility that the Sakhalin populations of *B. oreas* may also have been introduced. However, this hypothesis is unlikely for several reasons. First, all Sakhalin haplotypes were unique and not shared with Hokkaido haplotypes, indicating that the populations are native. Second, high genetic diversity was observed: nine haplotypes were detected from 13 specimens across five sites in Sakhalin. This result is difficult to explain by artificial introduction unless multiple introductions from genetically distinct source populations are assumed. Conversely, the Honshu populations, which are considered to have been artificially introduced, exhibited only two haplotypes shared with Southwestern Hokkaido populations. This pattern is commonly observed in artificially introduced populations (e.g., Machida et al., 2021; Okada et al., 2024). The genetic pattern in Sakhalin clearly differs from this. Finally, artificial introductions like those reported in previous studies are less likely to occur due to ecological differences between *B. oreas* and the two introduced loach species. While the two loach species prefer lentic environments such as slow-flowing streams or marshes, *B. oreas* primarily inhabits lotic river environments (Nakajima & Uchiyama, 2017), making unintentional co-introduction less likely.

This study revealed the phylogeographic structure of *B. oreas* and a notable pattern of back dispersal from Hokkaido to Sakhalin, challenging the conventional view of unidirectional colonization from the continent. Back dispersal has been widely reported in terrestrial taxa such as birds, insects, and plants (Heaney, 2007; Bellemain & Ricklefs, 2008; Patiño et al., 2017), but remains rarely documented in primary freshwater fishes, likely due to their lack of ability to disperse across marine barriers, making this case particularly noteworthy. Together with the remarkable northern-southern genetic structure and mixed ancestry patterns among neighboring subclusters within Hokkaido, our results suggest that repeated dispersal and isolation events, made possible by sea-level fluctuations during glacial-interglacial cycles in the Quaternary, have shaped the evolutionary history of this species. Further comparative phylogeographic studies across multiple taxa in Northeast Asia may provide deeper insights into the contribution of island systems in shaping continental biodiversity.

## Acknowledgments

We express our thanks Dr. T. Ito (Hokkaido Aquatic Biology), Prof. J. Miyazaki (Yamanashi University), Prof. K. Hosoya (Kinki University), Dr. A. Ohkawa (Tokyo University), Dr. Y. Takashima (Furano, Hokkaido), Dr. T. Suzuki (Hiroshima Shudo University), Dr. S. Komaki (Tohoku University), Dr. K. Yano (Lake Suwa Environmental Research Center), and many members in the Tojo Laboratory (Shinshu University), for valuable advice and encouragement for supporting our research. Work SVS and TSV was carried out within the assignment of Ministry of Science and Higher Education of the Russian Federation (theme No. 124012200182-1). This study was supported by KAKENHI (JSPS, Grants-in-Aid for Challenging Exploratory Research from the Japan Society for Promotion of Science; #23657064, KT), by grants for the River Fund of the River Foundation (#24-1215-016, #26-1215-017, KT), by a research grant from the Institute of Mountain Science, Shinshu University (KT).

## Author Contributions

KK, SVS, TSV and KT conducted sampling, and HN and KK conducted molecular experiments under the supervision of KT. HN and KK performed molecular phylogenetic analyses of the datasets using Sanger sequencing, and HN performed genome-wide SNPs analyses with the cooperation of MT, GU, and KT. HN prepared the initial draft, and completed the manuscript with significant input from all authors.

## Data Accessibility Statement

The DNA sequences have been deposited in the GenBank. All relevant methodological details are provided in this paper.

