## Supplementary Information for "Genetic structures of the Japanese stone loach *Barbatula oreas* (Cypriniformes: Nemacheilidae) in Sakhalin and Hokkaido: back dispersal from Hokkaido to Sakhalin"

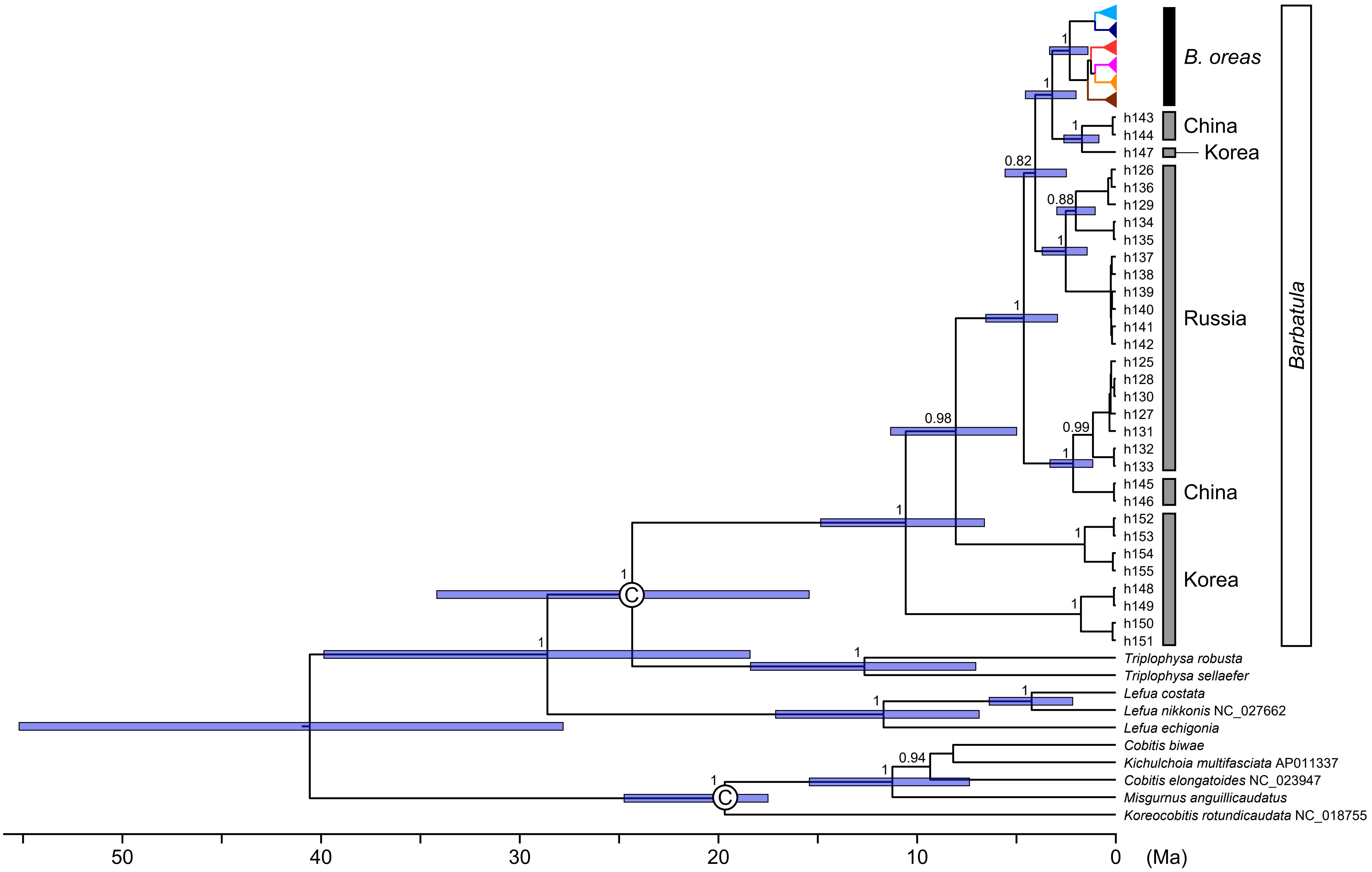


**Fig. S1.** Bayesian tree and calibration point based on the mtDNA *cytb* and 12S rRNA sequences.

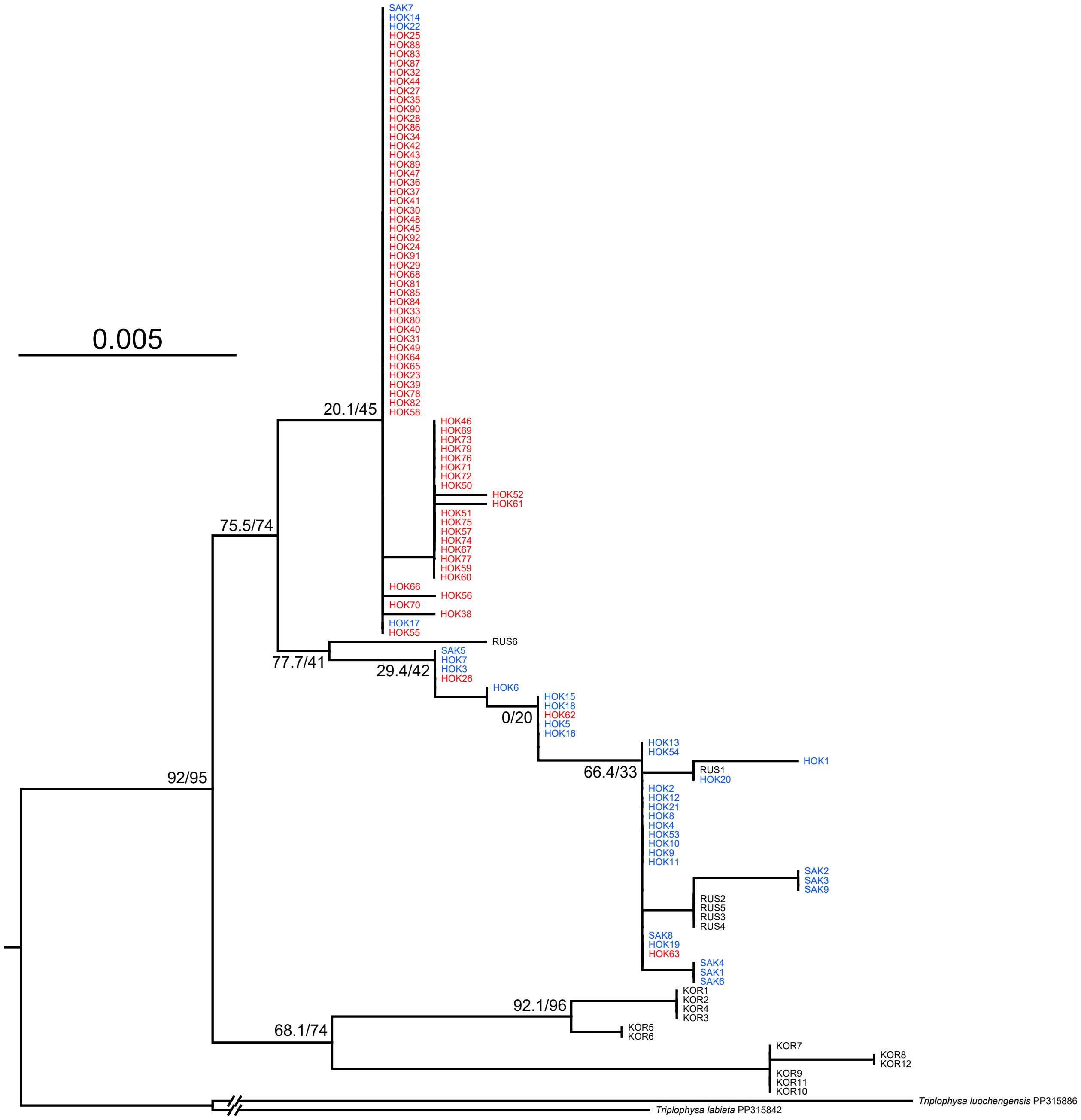


**Fig. S2.** Maximum likelihood tree of the genus *Barbatula* and the related species based on the nDNA *RAG1* (862 bp) sequences. The numbers at major nodes indicate SH-aLRT (left) and UFBoot (right) values. Specimens shown in blue correspond to the mtDNA Northern Clade, whereas those shown in red correspond to the mtDNA Southern Clade.

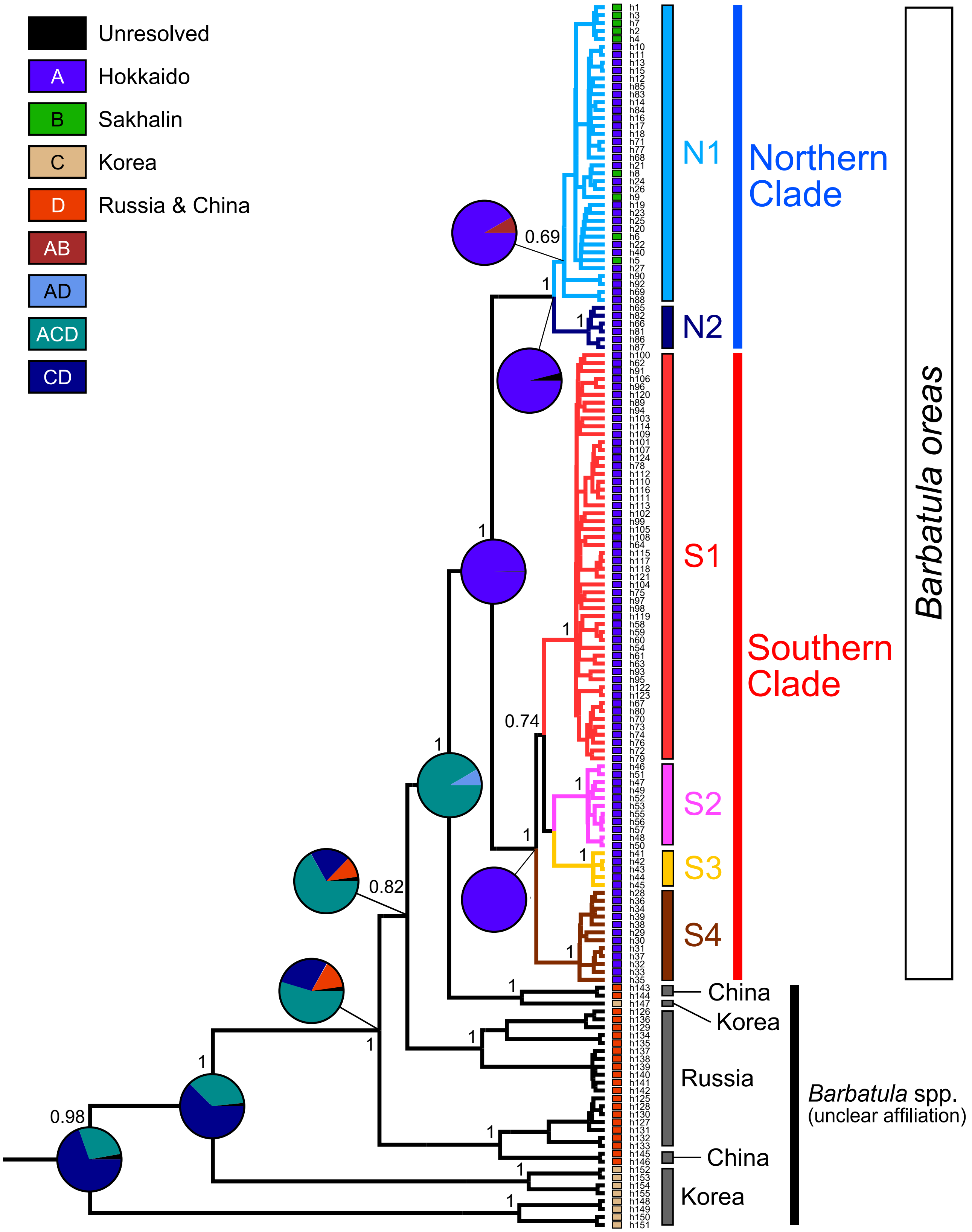


**Fig. S3.** Ancestral area reconstructions based on the Statistical Dispersal–Extinction–Cladogenesis (S-DEC) method.

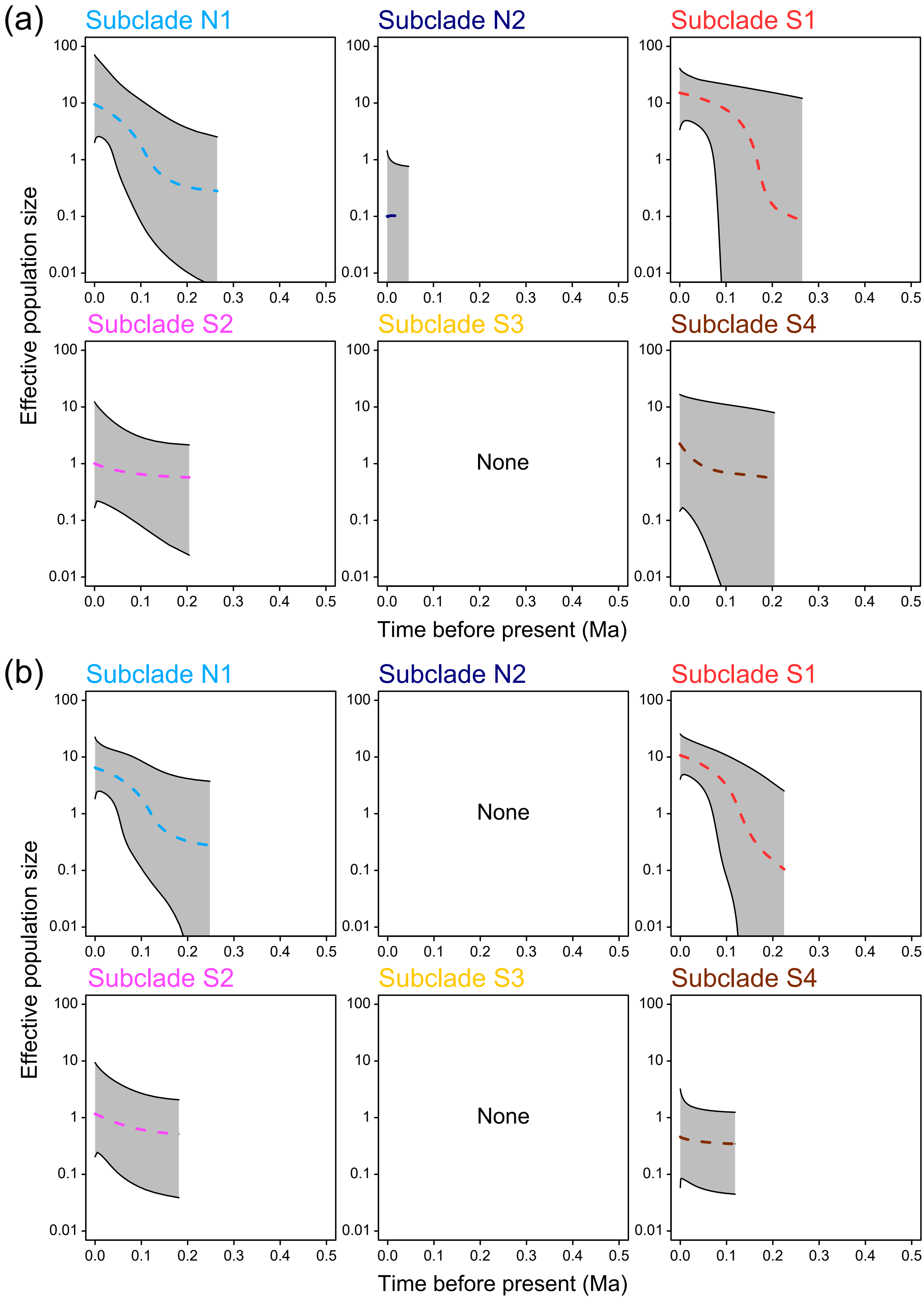


**Fig. S4.** Extended Bayesian Skyline Plot (EBSP) based on (a) the mtDNA 12S rRNA sequences, and (b) both mtDNA *cytb* and 12S rRNA sequences.

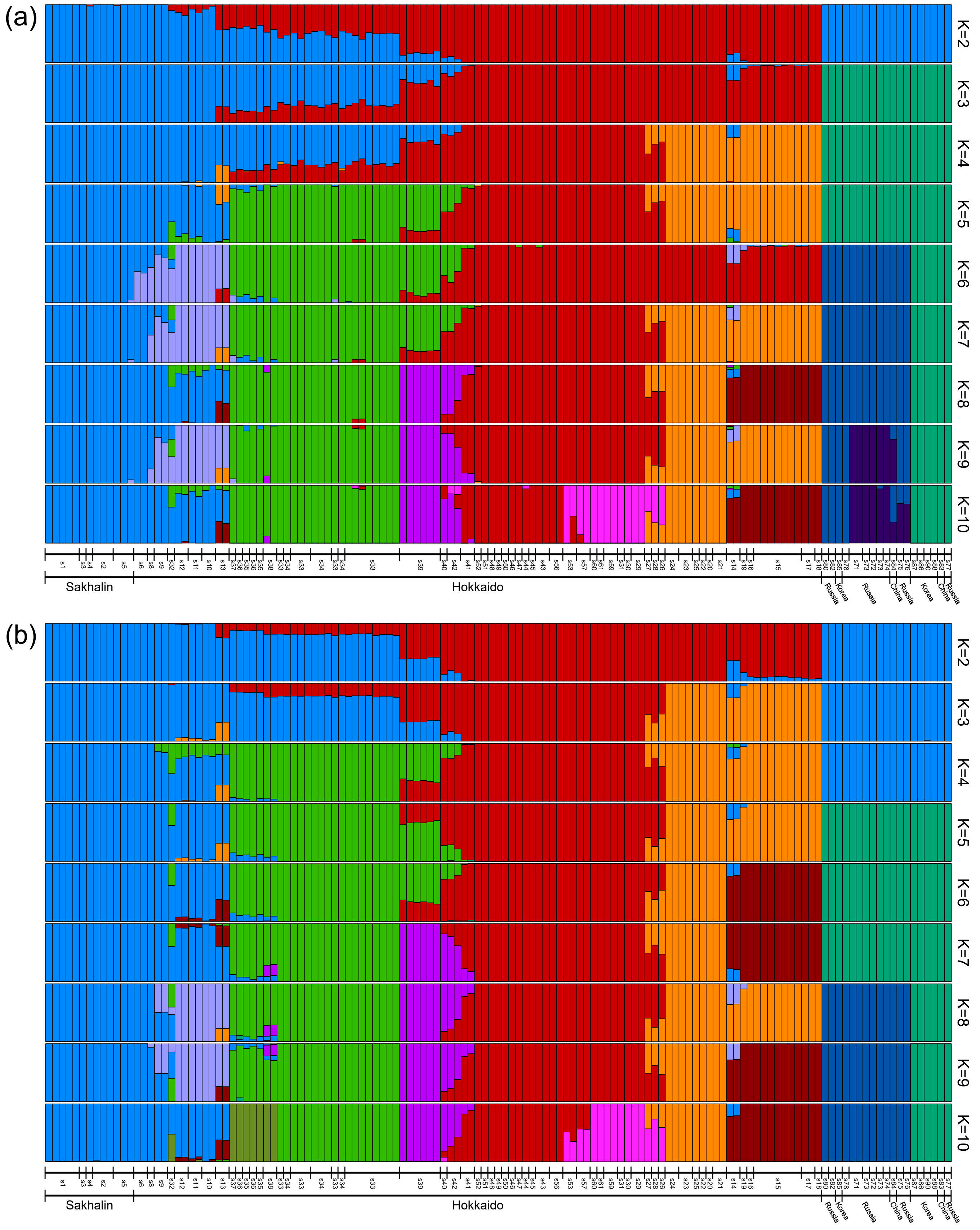


**Fig. S5.** ADMIXTURE plots for K = 2–10 of the genus *Barbatula* based on (a) *de novo* dataset 2 (2,178 SNPs) and (b) mapping dataset 3 (19,243 SNPs).

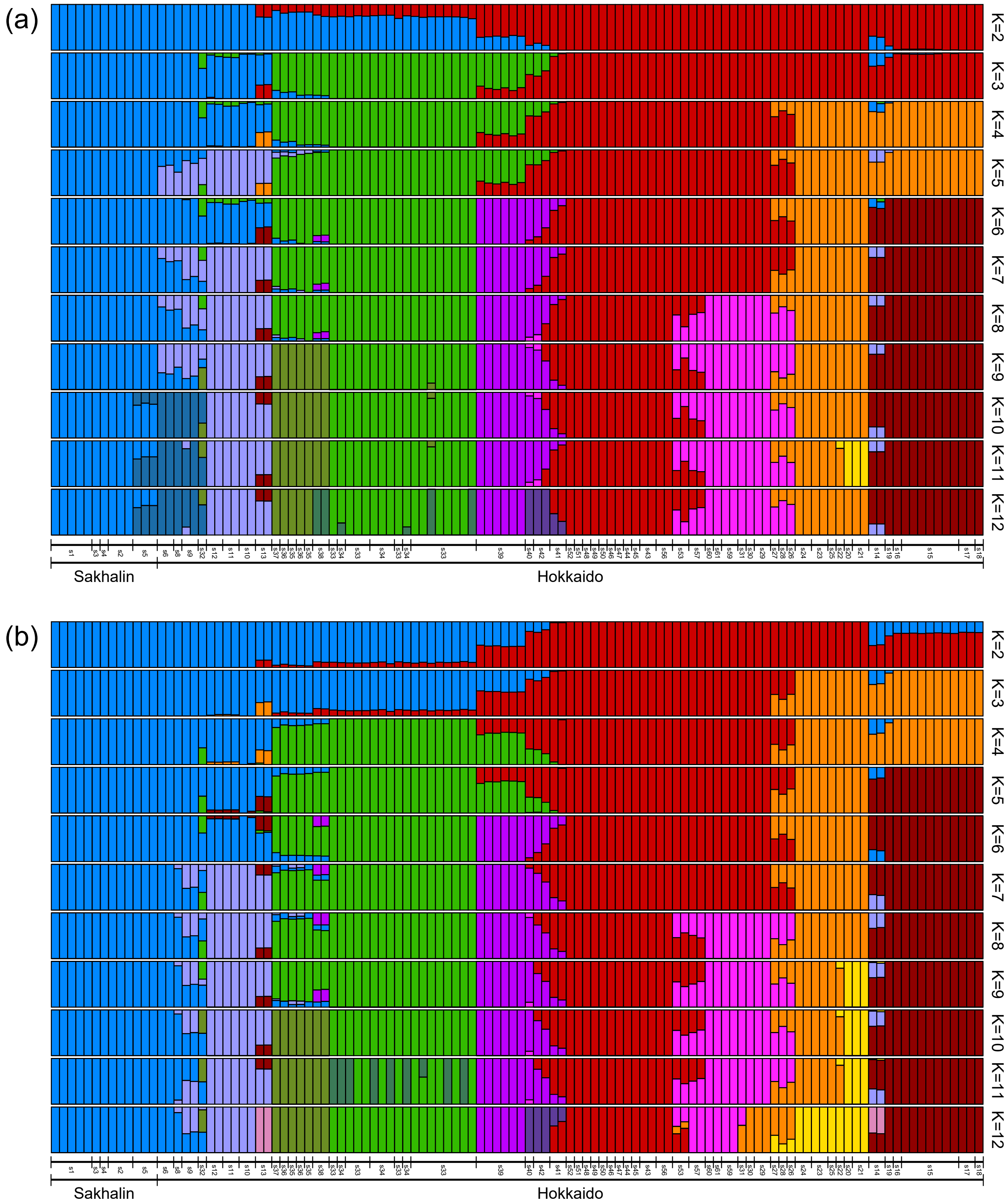


**Fig. S6.** ADMIXTURE plots for K = 2–12 of *Barbatula oreas* in Hokkaido and Sakhalin based on (a) *de novo* dataset 3 (8,178 SNPs) and (b) mapping dataset 7 (13,125 SNPs).

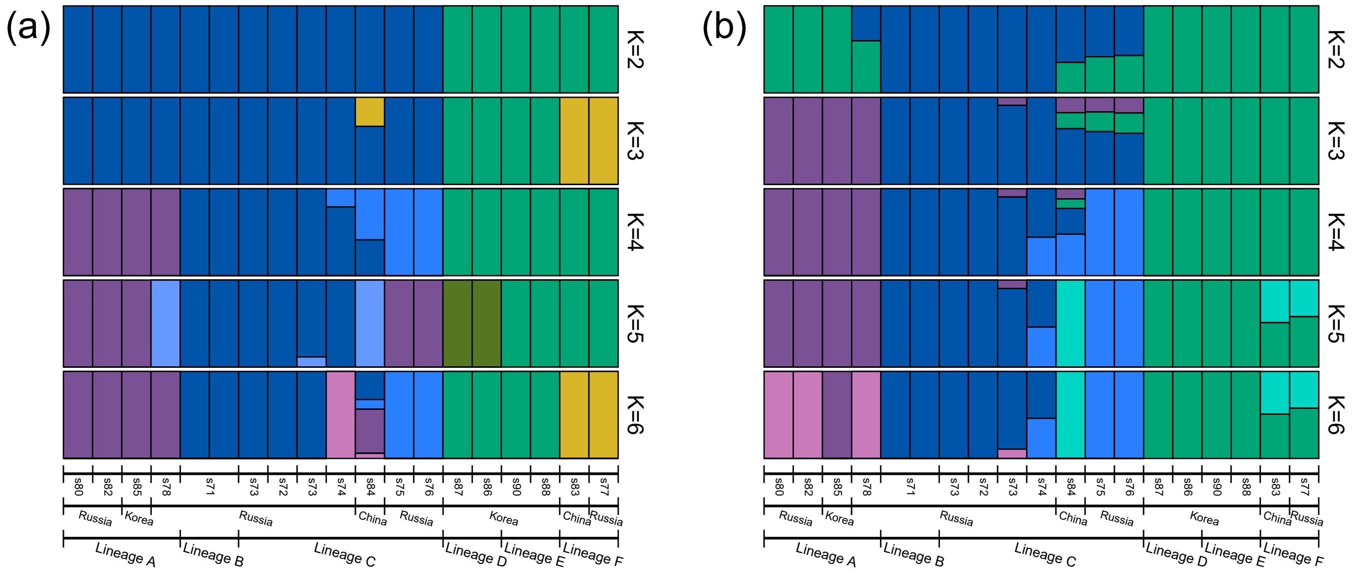


**Fig. S7.** ADMIXTURE plots for K = 2–6 of the genus *Barbatula* in continental East Asia based on (a) *de novo* dataset 5 (2,116 SNPs) and (b) mapping dataset 9 (7,247 SNPs).

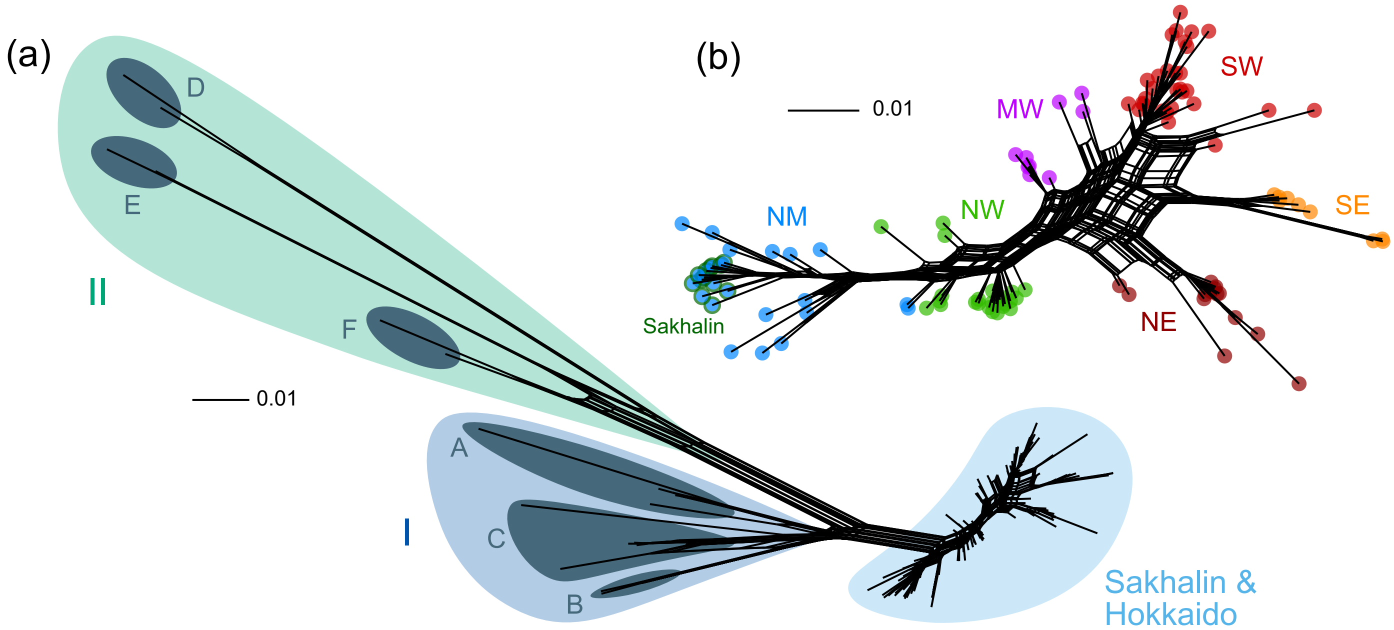


**Fig. S8.** Phylogenetic networks constructed using the Neighbor-net algorithm based on (a) *de novo* dataset 1 (all samples; 3,097 SNPs) and (b) *de novo* dataset 3 (*B. oreas* only; 8,178 SNPs).

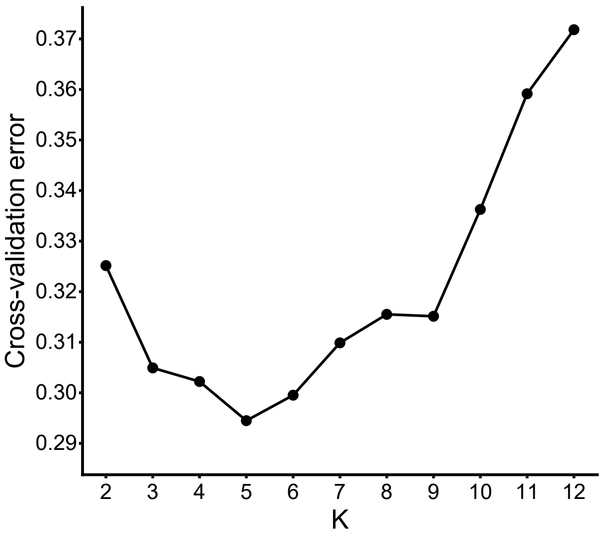


**Fig. S9.** Cross-validation error from ADMIXTURE analysis based on mapping dataset 7 (*B. oreas* only; 13,125 SNPs).

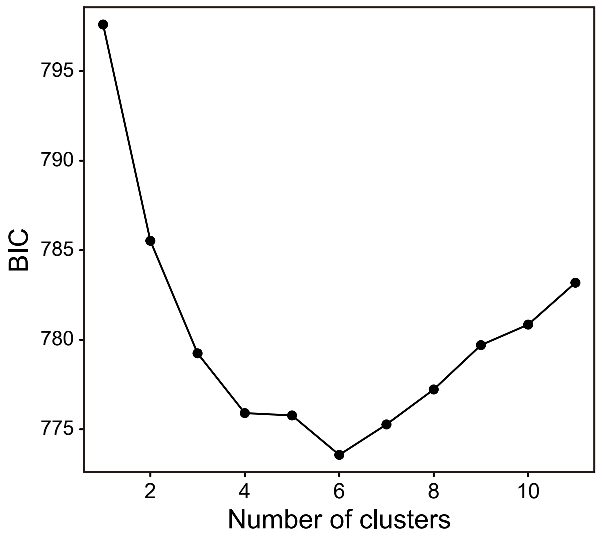


**Fig. S10.** BIC values from DAPC analysis based on mapping dataset 7 (*B. oreas* only; 13,125 SNPs).

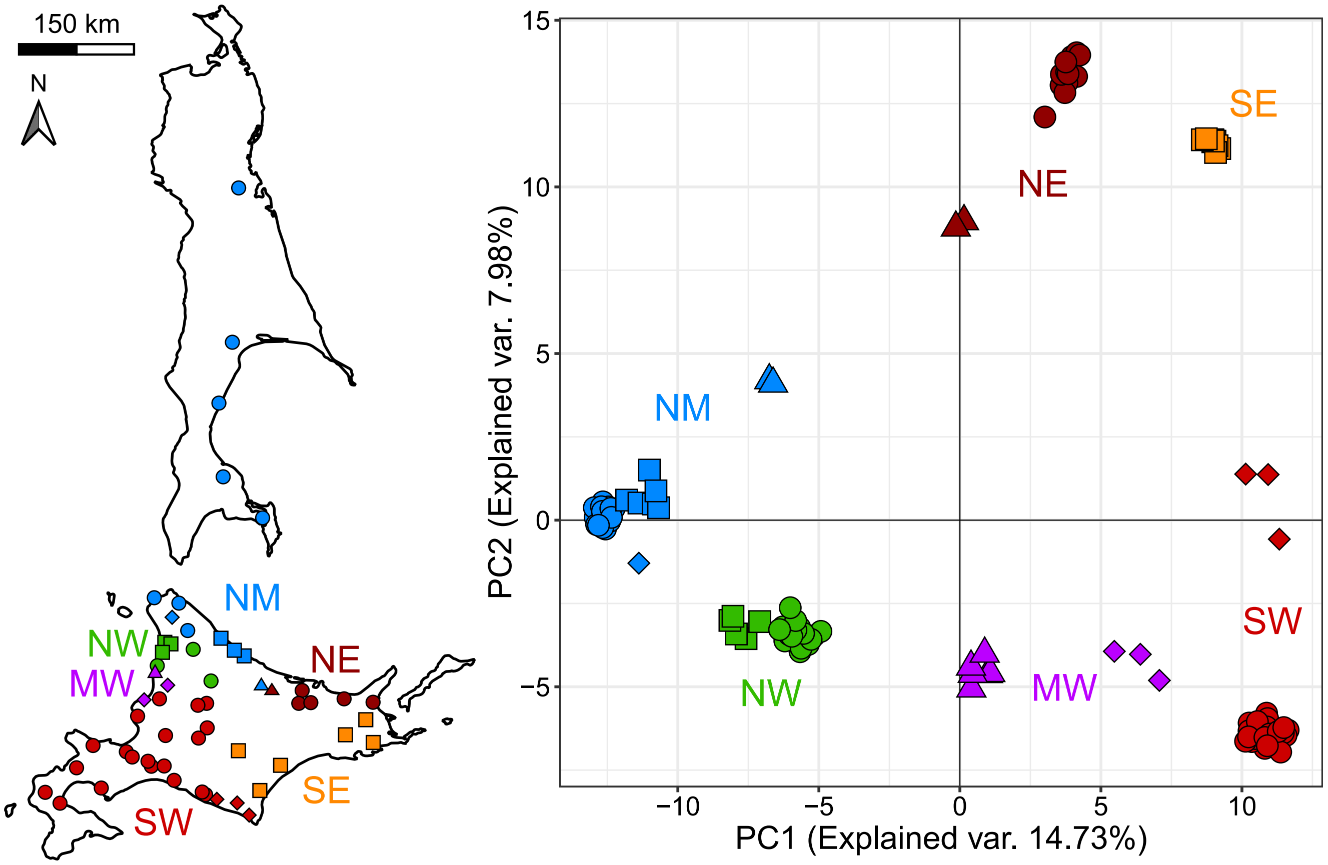


**Fig. S11.** PCA plot based on 13,125 nuclear SNPs. The colors and symbols correspond to the map on the left.

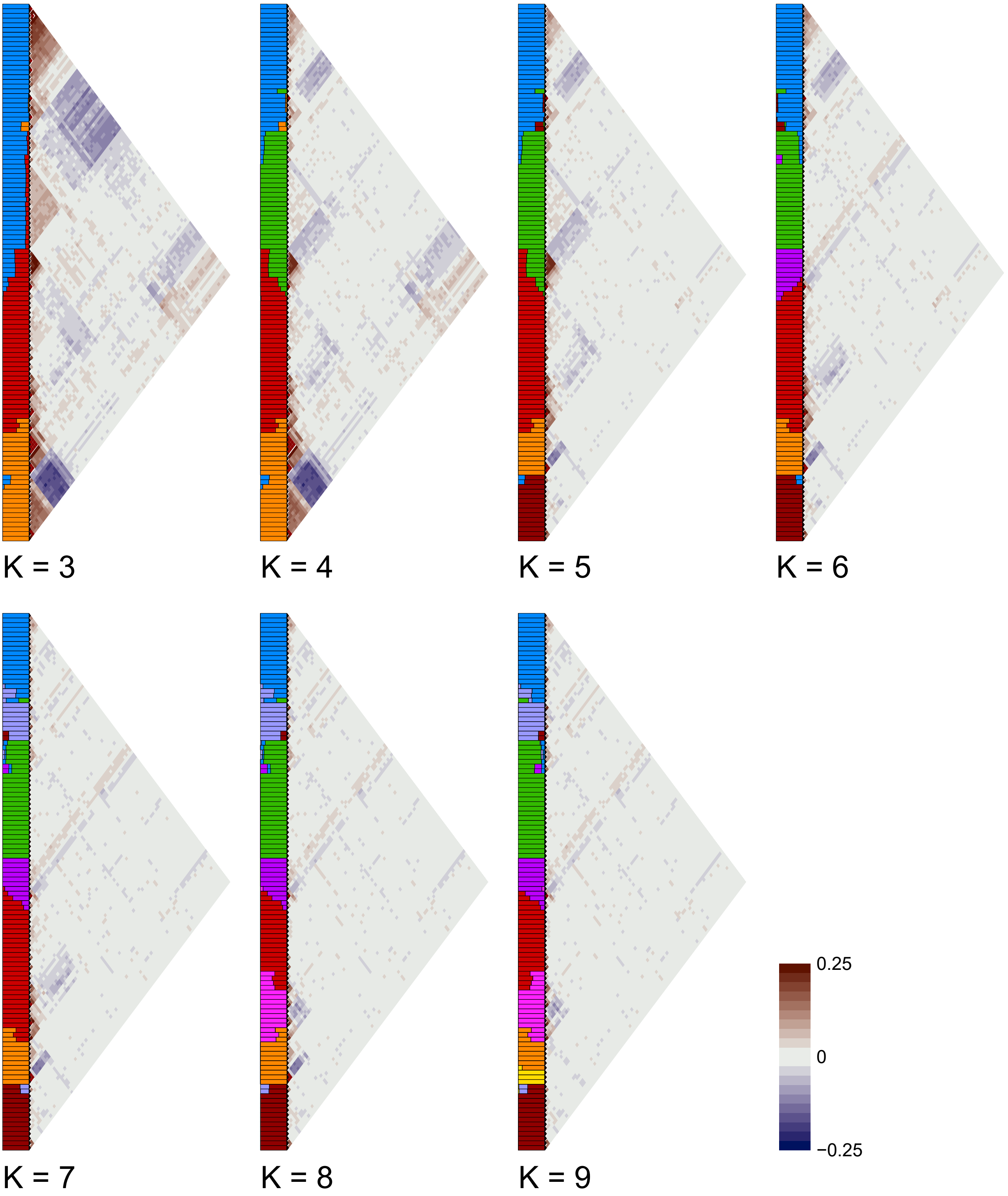


**Fig. S12.** Correlations of residuals between individuals to evaluate model fit using evalAdmix analysis based on ADMIXTURE results for K = 3–9 inferred from 13,125 nuclear SNPs. Positive (brown) and negative (blue) correlation values indicate poor fit to the inferred admixture model, whereas values close to zero indicate a good fit.

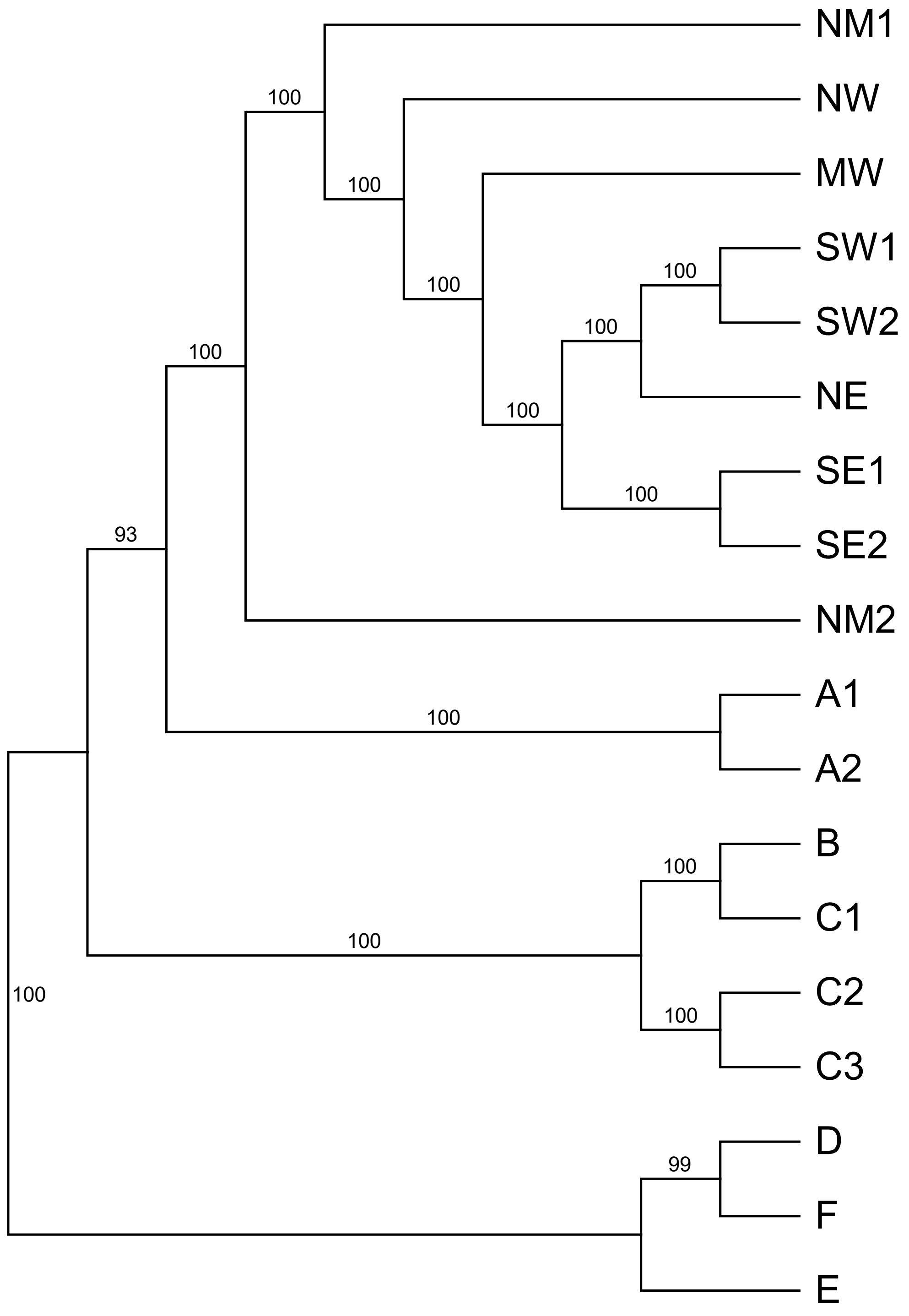


**Fig. S13.** Phylogenetic tree inferred using the SVDquartets method based on 159,567 SNPs. Bootstrap values are shown at major nodes.

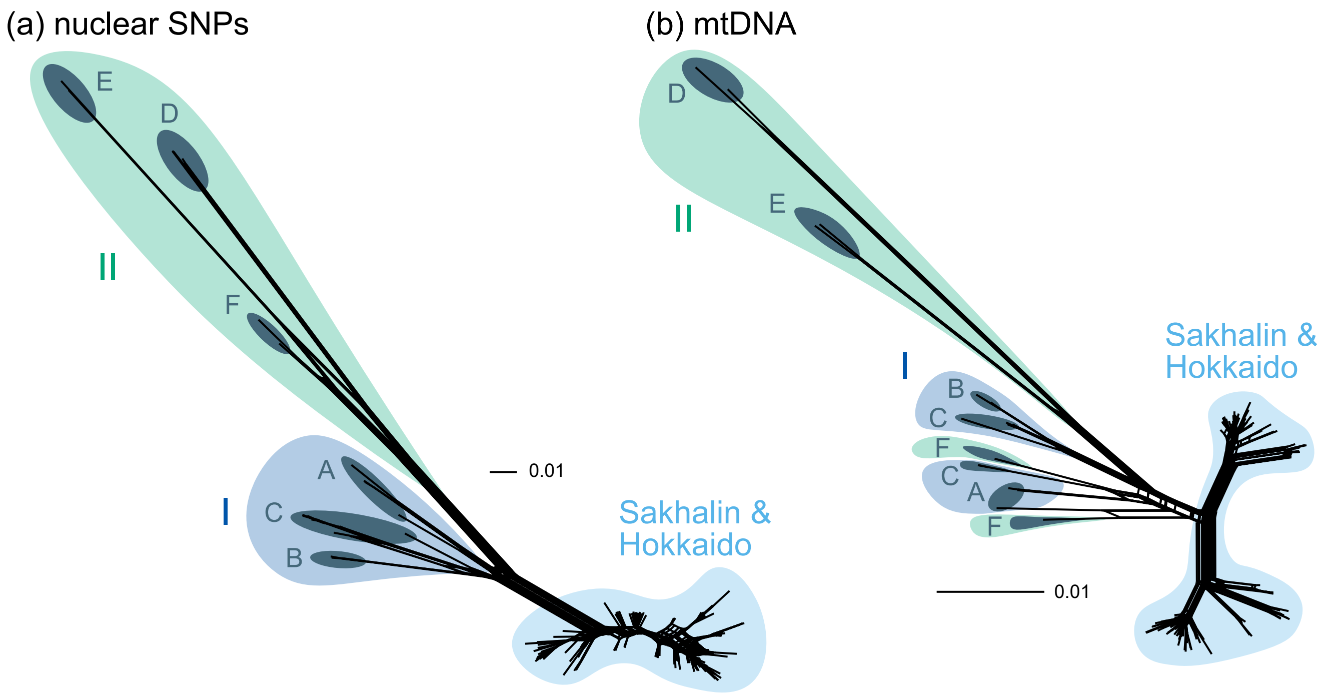


**Fig. S14.** Mito-nuclear discordance in Lineage F. Phylogenetic networks constructed using the Neighbor-net algorithm based on (a) nuclear SNPs (mapping dataset 1) and (b) six mtDNA sequences.

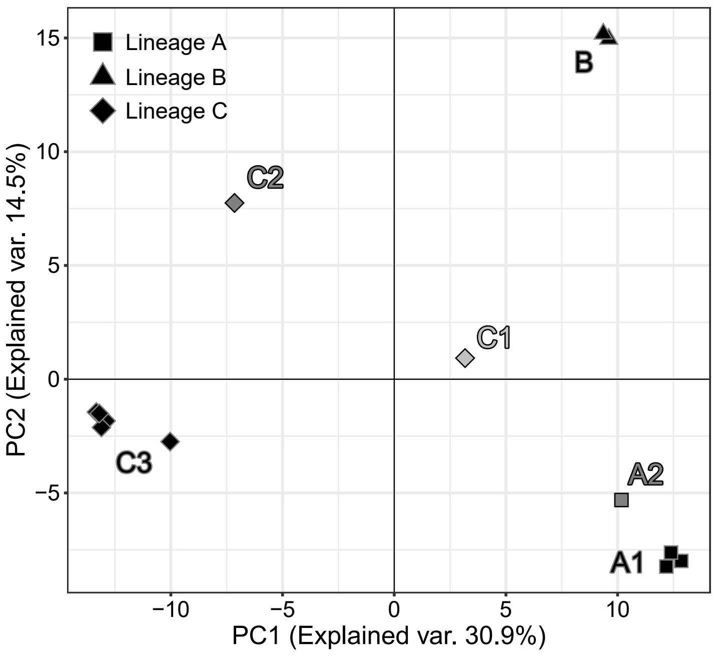


**Fig. S15.** PCA plot based on mapping dataset 10 (Lineages A–C only; 2,956 SNPs).

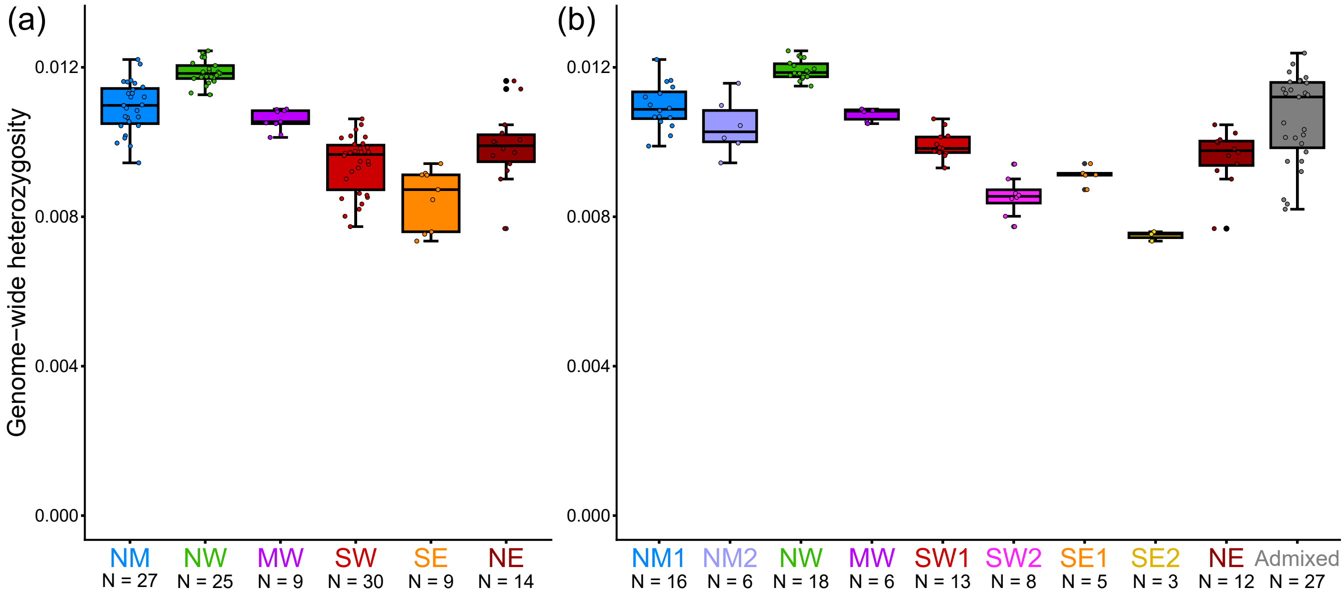


**Fig. S16.** Boxplots showing genome-wide heterozygosity for (a) major clusters and (b) subclusters.

### **Table S1.** List of specimens, sampling locations, mitochondrial haplotypes, and GenBank accession numbers of *Barbatula oreas*

| Site No. | Locality | Latitude | Longitude | Subclade | mtDNA Haplotype | | | nDNA *RAG1* OTU | SNPs OTU | SNPs Group | | Acces. No. | | | | | |
|---|---|---|---|---|---|---|---|---|---|---|---|---|---|---|---|---|---|
| | | | | | *cytb* | *cytb* + 12S | All mtDNA regions | | | Cluster | Subcluster | mtDNA cyt *b* | mtDNA *COI* | mtDNA D-loop | mtDNA 12S rRNA, tRNA-Val, and 16S rRNA | nDNA *RAG1* | SNPs |
| 1 | Tvm river, Sakhalin, Russia | 51.649 | 142.971 | N1 | n1 | h1 | H1 | SAK1 | s1_SAK_NOG1 | NM | NM1 | | | | | | |
| | | | | N1 | n2 | h2 | H2 | SAK2 | s1_SAK_NOG5 | NM | NM1 | | | | | | |
| | | | | N1 | n3 | h3 | H3 | SAK3 | s1_SAK_NOG4 | NM | NM1 | | | | | | |
| | | | | N1 | n4 | h4 | H4 | SAK4 | s1_SAK_NOG3 | NM | NM1 | | | | | | |
| | | | | N1 | n3 | h3 | H3 | SAK5 | s1_SAK_NOG2 | NM | NM1 | | | | | | |
| 2 | Maiskoe, Sakhalin, Russia | 49.293 | 142.875 | N1 | n5 | h5 | H5 | - | s2_SAK_MAI1 | NM | NM1 | | | | | | |
| | | | | N1 | n5 | h5 | H6 | SAK6 | s2_SAK_MAI2 | NM | NM1 | | | | | | |
| | | | | N1 | n1 | h1 | H7 | SAK7 | s2_SAK_MAI3 | NM | NM1 | | | | | | |
| 3 | Zaozernoe, Sakhalin, Russia | 48.364 | 142.670 | N1 | n6 | h6 | H8 | SAK8 | s3_SAK_ZAO | NM | NM1 | | | | | | |
| 4 | Sokol, Sakhalin, Russia | 47.239 | 142.735 | N1 | n7 | h7 | H9 | - | s4_SAK_SOK | NM | NM1 | | | | | | |
| 5 | Pikhtovoe, Sakhalin, Russia | 46.612 | 143.337 | N1 | n8 | h8 | H10 | - | s5_SAK_PIK1 | NM | NM1 | | | | | | |
| | | | | N1 | n9 | h9 | H11 | SAK9 | s5_SAK_PIK2 | NM | NM1 | | | | | | |
| | | | | N1 | n1 | h1 | H8 | - | s5_SAK_PIK3 | NM | NM1 | | | | | | |
| 6 | Midori, Wakkanai, Hokkaido | 45.393 | 141.683 | N1 | n10 | h10 | H12 | HOK1 | s6_HOK_WAK1 | NM | NM1 | | | | | | |
| | | | | N1 | n10 | h11 | H13 | HOK2 | - | - | - | | | | | | |
| | | | | N1 | n10 | h10 | H12 | HOK3 | - | - | - | | | | | | |
| | | | | N1 | n11 | h12 | H14 | - | s6_HOK_WAK2 | NM | NM1 | | | | | | |
| 7 | Soya, Wakkanai, Hokkaido | 45.334 | 141.866 | N1 | n12 | h13 | H15 | HOK4 | - | - | - | | | | | | |
| | | | | N1 | n13 | h14 | H16 | HOK5 | - | - | - | | | | | | |
| | | | | N1 | n12 | h15 | H17 | HOK6 | - | - | - | | | | | | |
| | | | | N1 | n12 | h13 | H18 | HOK7 | - | - | - | | | | | | |
| 8 | Sarufutsu, Hokkaido | 45.310 | 142.059 | N1 | n14 | h16 | H19 | HOK8 | s8_HOK_SAR | NM | NM1 | | | | | | |
| | | | | N1 | n13 | h14 | H20 | HOK9 | - | - | - | | | | | | |
| | | | | N1 | n13 | h14 | H20 | HOK10 | - | - | - | | | | | | |
| | | | | N1 | n14 | h16 | H19 | HOK11 | - | - | - | | | | | | |
| 9 | Nakatombetsu, Hokkaido | 44.892 | 142.194 | N1 | n13 | h14 | H21 | HOK12 | s9_HOK_NAK1 | NM | Admixed | | | | | | |
| | | | | N1 | n15 | h17 | H22 | HOK13 | s9_HOK_NAK2 | NM | Admixed | | | | | | |
| | | | | N1 | n13 | h14 | H20 | HOK14 | - | - | - | | | | | | |
| | | | | N1 | n16 | h18 | H23 | HOK15 | - | - | - | | | | | | |
| 10 | Esashi, Hokkaido | 44.774 | 142.701 | N1 | n17 | h19 | H24 | - | s10_HOK_ESA1 | NM | NM2 | | | | | | |
| | | | | N1 | n17 | h20 | H25 | - | s10_HOK_ESA2 | NM | NM2 | | | | | | |
| | | | | N1 | n17 | h19 | H26 | - | - | - | - | | | | | | |
| | | | | N1 | n17 | h20 | H25 | - | - | - | - | | | | | | |
| 11 | Kitaomu, Omu, Hokkaido | 44.591 | 142.906 | N1 | n17 | h20 | H25 | HOK16 | s11_HOK_OMU_KIT1 | NM | NM2 | | | | | | |
| | | | | N1 | n18 | h21 | H27 | HOK17 | s11_HOK_OMU_KIT2 | NM | NM2 | | | | | | |
| | | | | N1 | n17 | h20 | H25 | HOK18 | - | - | - | | | | | | |
| | | | | N1 | n17 | h20 | H25 | HOK19 | - | - | - | | | | | | |
| 12 | Kamisawaki, Omu, Hokkaido | 44.505 | 143.056 | N1 | n17 | h19 | H28 | HOK20 | s12_HOK_OMU_KAM1 | NM | NM2 | | | | | | |
| | | | | N1 | n17 | h19 | H28 | HOK21 | - | - | - | | | | | | |
| | | | | N1 | n19 | h21 | H29 | HOK22 | s12_HOK_OMU_KAM2 | NM | NM2 | | | | | | |
| 13 | Kamikonomai, Monbetsu, Hokkaido | 44.093 | 143.317 | N1 | n17 | h19 | H30 | - | s13_HOK_MON1 | NM | Admixed | | | | | | |
| | | | | N1 | n20 | h23 | H31 | - | - | - | - | | | | | | |
| | | | | N1 | n17 | h19 | H30 | - | - | - | - | | | | | | |
| | | | | N1 | n21 | h24 | H32 | - | s13_HOK_MON2 | NM | Admixed | | | | | | |

| | | | | | | | | | | | |
|---|---|---|---|---|---|---|---|---|---|---|---|
| 14 | Engaru, Hokkaido | 44.021 | 143.478 | N1 | n17 | h19 | H33 | - | - | - | - |
| | | | | N1 | n17 | h19 | H34 | - | - | - | - |
| | | | | N1 | n22 | h25 | H35 | - | s14_HOK_ENG2 | NE | Admixed |
| | | | | N1 | n23 | h26 | H36 | - | s14_HOK_ENG1 | NE | Admixed |
| 15 | Tokoro, Kitami, Hokkaido | 43.977 | 143.941 | N1 | n17 | h27 | H37 | - | s15_HOK_KIT_TOK3 | NE | NE |
| | | | | S4 | s1 | h28 | H38 | - | s15_HOK_KIT_TOK1 | NE | NE |
| | | | | N1 | n17 | h27 | H37 | - | s15_HOK_KIT_TOK2 | NE | NE |
| | | | | S4 | s1 | h28 | H39 | - | s15_HOK_KIT_TOK5 | NE | NE |
| | | | | S4 | s1 | h28 | H38 | - | s15_HOK_KIT_TOK7 | NE | NE |
| | | | | S4 | s2 | h29 | H40 | - | - | - | - |
| | | | | N1 | n17 | h19 | H33 | - | s15_HOK_KIT_TOK6 | NE | NE |
| | | | | S4 | s3 | h30 | H41 | - | s15_HOK_KIT_TOK4 | NE | NE |
| | | | | S4 | s4 | h31 | H42 | - | - | - | - |
| | | | | S4 | s5 | h32 | H43 | - | - | - | - |
| 16 | Minamioka, Kitami, Hokkaido | 43.780 | 143.889 | S4 | s6 | h33 | H44 | HOK23 | s16_HOK_KIT_MIN | NE | NE |
| | | | | S4 | s2 | h29 | H45 | HOK24 | - | - | - |
| | | | | S4 | s7 | h34 | H46 | HOK25 | - | - | - |
| | | | | S4 | s8 | h35 | H47 | HOK26 | - | - | - |
| 17 | Bihoro, Hokkaido | 43.791 | 144.073 | S4 | s1 | h28 | H48 | - | s17_HOK_BIH2 | NE | NE |
| | | | | S4 | s1 | h36 | H49 | - | - | - | - |
| | | | | S4 | s9 | h37 | H50 | - | s17_HOK_BIH1 | NE | NE |
| | | | | S4 | s10 | h38 | H51 | - | - | - | - |
| 18 | Kiyosato, Hokkaido | 43.849 | 144.581 | S4 | s1 | h39 | H52 | - | s18_HOK_KIY | NE | NE |
| | | | | S4 | s1 | h39 | H52 | - | - | - | - |
| | | | | S4 | s1 | h39 | H52 | - | - | - | - |
| | | | | S4 | s1 | h39 | H52 | - | - | - | - |
| | | | | S4 | s1 | h39 | H52 | - | - | - | - |
| 19 | Kunbetsu, Shibetsu, Hokkaido | 43.797 | 145.023 | N1 | n24 | h40 | H53 | - | s19_HOK_SHI_KUN | NE | NE |
| | | | | N1 | n24 | h40 | H53 | - | - | - | - |
| | | | | N1 | n24 | h40 | H53 | - | - | - | - |
| 20 | Aobadai, Nakashibetsu, Hokkaido | 43.530 | 144.912 | S3 | s11 | h41 | H54 | HOK27 | s20_HOK_NAK | SE | SE2 |
| | | | | S3 | s12 | h42 | H55 | HOK28 | - | - | - |
| | | | | S3 | s13 | h43 | H56 | HOK29 | - | - | - |
| | | | | S3 | s11 | h41 | H57 | HOK30 | - | - | - |
| 21 | Hamanaka, Hokkaido | 43.182 | 145.026 | S3 | s11 | h41 | H58 | HOK31 | s21_HOK_HAM1 | SE | SE2 |
| | | | | S3 | s14 | h44 | H59 | HOK32 | s21_HOK_HAM2 | SE | SE2 |
| | | | | S3 | s11 | h41 | H60 | HOK33 | - | - | - |
| | | | | S3 | s15 | h45 | H61 | HOK34 | - | - | - |
| 22 | Shibecha, Hokkaido | 43.300 | 144.602 | S2 | s16 | h46 | H62 | HOK35 | s22_HOK_SHI_KAI | SE | Admixed |
| | | | | S2 | s17 | h47 | H63 | HOK36 | - | - | - |
| | | | | S2 | s16 | h46 | H64 | - | - | - | - |
| | | | | S2 | s16 | h46 | H64 | HOK37 | - | - | - |
| 23 | Urahoro, Hokkaido | 42.834 | 143.611 | S2 | s18 | h48 | H65 | HOK38 | s23_HOK_URA1 | SE | SE1 |
| | | | | S2 | s17 | h49 | H66 | HOK39 | - | - | - |
| | | | | S2 | s18 | h50 | H67 | HOK40 | - | - | - |
| | | | | S2 | s19 | h51 | H68 | HOK41 | s23_HOK_URA2 | SE | SE1 |
| 24 | Shimizu, Hokkaido | 43.057 | 142.968 | S2 | s17 | h49 | H69 | - | s24_HOK_SHI_BIB1 | SE | SE1 |
| | | | | S2 | s19 | h51 | H70 | - | s24_HOK_SHI_BIB2 | SE | SE1 |
| | | | | S2 | s17 | h49 | H69 | - | - | - | - |

| | | | | | | | | | | | |
|---|---|---|---|---|---|---|---|---|---|---|---|
| 25 | Taiki, Hokkaido | 42.449 | 143.294 | S2 | s20 | h52 | H71 | - | s25_HOK_TAI | SE | SE1 |
| | | | | S2 | s17 | h49 | H72 | - | - | - | - |
| | | | | S2 | s17 | h49 | H73 | - | - | - | - |
| | | | | S2 | s17 | h49 | H72 | - | - | - | - |
| 26 | Erimo, Hokkaido | 42.074 | 143.132 | S2 | s21 | h53 | H74 | HOK42 | s26_HOK_ERI | SW | Admixed |
| | | | | S2 | s21 | h53 | H74 | HOK43 | - | - | - |
| | | | | S2 | s21 | h53 | H74 | HOK44 | - | - | - |
| | | | | S2 | s21 | h53 | H74 | HOK45 | - | - | - |
| 27 | Urakawa, Hokkaido | 42.259 | 142.949 | S1 | s22 | h54 | H75 | HOK46 | s27_HOK_URA | SW | Admixed |
| | | | | S2 | s23 | h55 | H76 | HOK47 | - | - | - |
| | | | | S2 | s23 | h56 | H77 | HOK48 | - | - | - |
| | | | | S2 | s23 | h57 | H78 | HOK49 | - | - | - |
| 28 | Mitsuishi, Shinhidaka, Hokkaido | 42.315 | 142.638 | S2 | s21 | h53 | H79 | - | s28_HOK_SHI_MIT | SW | Admixed |
| | | | | S2 | s21 | h53 | H79 | - | - | - | - |
| | | | | S2 | s21 | h53 | H79 | - | - | - | - |
| | | | | S2 | s21 | h53 | H79 | - | - | - | - |
| 29 | Shizunai, Shinhidaka, Hokkaido | 42.387 | 142.467 | S1 | s22 | h58 | H80 | HOK50 | s29_HOK_SHI_SHI2 | SW | SW2 |
| | | | | S1 | s24 | h59 | H81 | HOK51 | s29_HOK_SHI_SHI1 | SW | SW2 |
| | | | | S1 | s25 | h60 | H82 | HOK52 | - | - | - |
| 30 | Niikappu, Hokkaido | 42.426 | 142.415 | S1 | s22 | h58 | H83 | - | s30_HOK_NII | SW | SW2 |
| | | | | S1 | s22 | h58 | H84 | - | - | - | - |
| 31 | Mukawa, Hokkaio | 42.605 | 141.986 | S1 | s25 | h61 | H85 | - | s31_HOK_MUK | SW | SW2 |
| | | | | S1 | s26 | h62 | H86 | - | - | - | - |
| | | | | S1 | s22 | h63 | H87 | - | - | - | - |
| | | | | S1 | s22 | h64 | H88 | - | - | - | - |
| 32 | Toyotomi, Hokkaido | 45.094 | 141.956 | N2 | n25 | h65 | H89 | HOK53 | s32_HOK_TOY | NM | Admixed |
| | | | | N2 | n25 | h65 | H90 | HOK54 | - | - | - |
| 33 | Bifuka, Hokkaido | 44.605 | 142.277 | N2 | n26 | h66 | H91 | - | s33_HOK_BIF11 | NW | NW |
| | | | | S1 | s27 | h67 | H92 | - | s33_HOK_BIF9 | NW | NW |
| | | | | N1 | n27 | h68 | H93 | - | s33_HOK_BIF13 | NW | NW |
| | | | | N2 | n26 | h66 | H91 | - | s33_HOK_BIF5 | NW | NW |
| | | | | N1 | n28 | h69 | H94 | - | s33_HOK_BIF3 | NW | NW |
| | | | | S1 | s28 | h70 | H95 | - | s33_HOK_BIF12 | NW | NW |
| | | | | N1 | n13 | h14 | H96 | - | s33_HOK_BIF1 | NW | NW |
| | | | | S1 | s22 | h64 | H97 | - | s33_HOK_BIF7 | NW | NW |
| | | | | N1 | n29 | h71 | H98 | - | s33_HOK_BIF2 | NW | NW |
| | | | | S1 | s29 | h72 | H99 | - | - | - | - |
| | | | | S1 | s30 | h73 | H100 | - | s33_HOK_BIF6 | NW | NW |
| | | | | S1 | s31 | h74 | H101 | - | - | - | - |
| | | | | S1 | s32 | h75 | H102 | - | s33_HOK_BIF4 | NW | NW |
| | | | | S1 | s33 | h76 | H103 | - | s33_HOK_BIF8 | NW | NW |
| | | | | N1 | n30 | h77 | H104 | - | s33_HOK_BIF10 | NW | NW |
| 34 | Kamishibetsu, Shibetsu, Hokkaido | 44.122 | 142.549 | S1 | s34 | h78 | H105 | - | s34_HOK_SHI3 | NW | NW |
| | | | | S1 | s35 | h79 | H106 | - | s34_HOK_SHI5 | NW | NW |
| | | | | S1 | s28 | h70 | H107 | - | s34_HOK_SHI2 | NW | NW |
| | | | | S1 | s36 | h80 | H108 | - | s34_HOK_SHI1 | NW | NW |
| | | | | N2 | n31 | h81 | H109 | - | s34_HOK_SHI4 | NW | NW |
| | | | | S1 | s28 | h70 | H95 | - | - | - | - |
| 35 | Kyuko, Embetsu, Hokkaido | 44.712 | 141.844 | N2 | n32 | h82 | H110 | - | s35_HOK_EMB_KYU1 | NW | Admixed |
| | | | | N1 | n13 | h83 | H111 | - | s35_HOK_EMB_KYU2 | NW | Admixed |

| | | | | | | | | | | | |
|---|---|---|---|---|---|---|---|---|---|---|---|
| 36 | Chuo, Embetsu, Hokkaido | 44.688 | 141.937 | N1 | n13 | h84 | H112 | - | s36_HOK_EMB_CHU1 | NW | Admixed |
| | | | | N2 | n26 | h66 | H113 | - | s36_HOK_EMB_CHU2 | NW | Admixed |
| | | | | N1 | n13 | h84 | H114 | - | - | - | - |
| | | | | N1 | n33 | h85 | H115 | - | - | - | - |
| 37 | Shosanbetsu, Hokkaido | 44.559 | 141.808 | N2 | n34 | h86 | H116 | - | s37_HOK_SHO | NW | Admixed |
| | | | | N2 | n35 | h87 | H117 | - | - | - | - |
| | | | | N2 | n34 | h86 | H116 | - | - | - | - |
| | | | | N2 | n35 | h87 | H118 | - | - | - | - |
| 38 | Haboro, Hokkaido | 44.355 | 141.729 | N1 | n13 | h84 | H119 | - | - | - | - |
| | | | | N2 | n34 | h86 | H120 | - | s38_HOK_HAB2 | NW | Admixed |
| | | | | N1 | n28 | h88 | H121 | - | s38_HOK_HAB1 | NW | Admixed |
| | | | | N1 | n28 | h69 | H122 | - | - | - | - |
| 39 | Tomamae, Hokkaido | 44.282 | 141.696 | S1 | s37 | h89 | H123 | - | s39_HOK_TOM2 | MW | MW |
| | | | | N1 | n36 | h90 | H124 | - | s39_HOK_TOM5 | MW | MW |
| | | | | S1 | s38 | h91 | H125 | - | s39_HOK_TOM1 | MW | MW |
| | | | | S1 | s38 | h91 | H125 | - | s39_HOK_TOM6 | MW | MW |
| | | | | N1 | n36 | h92 | H126 | - | s39_HOK_TOM4 | MW | MW |
| | | | | S1 | s37 | h89 | H123 | - | s39_HOK_TOM3 | MW | MW |
| 40 | Obira, Hokkaido | 44.057 | 141.888 | S1 | s39 | h93 | H127 | HOK55 | s40_HOK_OBI | MW | Admixed |
| | | | | S1 | s22 | h94 | H128 | HOK56 | - | - | - |
| 41 | Rumoi, Rumoi, Hokkaido | 43.849 | 141.763 | S1 | s34 | h78 | H129 | HOK57 | s41_HOK_RUM2 | SW | Admixed |
| | | | | S1 | s40 | h95 | H130 | HOK58 | s41_HOK_RUM1 | SW | Admixed |
| | | | | S1 | s40 | h95 | H130 | HOK59 | - | - | - |
| | | | | S1 | s41 | h96 | H131 | HOK60 | - | - | - |
| 42 | Mashike, Hokkaido | 43.834 | 141.529 | S1 | s22 | h64 | H132 | HOK61 | s42_HOK_MAS2 | MW | Admixed |
| | | | | S1 | s22 | h64 | H133 | HOK62 | - | - | - |
| | | | | S1 | s42 | h97 | H134 | HOK63 | s42_HOK_MAS1 | MW | Admixed |
| 43 | Hamamasu, Ishikari, Hokkaido | 43.585 | 141.429 | S1 | s22 | h64 | H135 | HOK64 | s43_HOK_ISH2 | SW | SW1 |
| | | | | S1 | s22 | h64 | H136 | HOK65 | s43_HOK_ISH1 | SW | SW1 |
| | | | | S1 | s22 | h64 | H137 | HOK66 | - | - | - |
| | | | | S1 | s22 | h64 | H138 | HOK67 | - | - | - |
| 44 | Higashiasahikawa, Asahikawa, Hokkaido | 43.780 | 142.483 | S1 | s22 | h64 | H139 | - | - | - | - |
| | | | | S1 | s43 | h98 | H140 | - | s44_HOK_ASA_HIG | SW | SW1 |
| | | | | S1 | s22 | h99 | H141 | - | - | - | - |
| | | | | S1 | s44 | h100 | H142 | - | - | - | - |
| 45 | Kagura, Asahikawa, Hokkaido | 43.751 | 142.349 | S1 | s43 | h98 | H140 | - | - | - | - |
| | | | | S1 | s43 | h98 | H143 | - | - | - | - |
| | | | | S1 | s34 | h78 | H129 | - | s45_HOK_ASA_KAG | SW | SW1 |
| 46 | Kamifurano, Hokkaido | 43.406 | 142.491 | S1 | s22 | h64 | H144 | HOK68 | s46_HOK_KAM | SW | SW1 |
| 47 | Furano, Hokkaido | 43.250 | 142.359 | S1 | s45 | h101 | H145 | HOK69 | s47_HOK_FUR | SW | SW1 |
| | | | | S1 | s46 | h102 | H146 | HOK70 | - | - | - |
| | | | | S1 | s22 | h64 | H147 | HOK71 | - | - | - |
| 48 | Koshunai, Bibai, Hokkaido | 43.289 | 141.855 | S1 | s22 | h64 | H144 | - | s48_HOK_BIB | SW | SW1 |
| | | | | S1 | s22 | h103 | H148 | - | - | - | - |
| 49 | Nishi, Sapporo, Hokkaido | 43.041 | 141.256 | S1 | s22 | h64 | H149 | HOK72 | s49_HOK_SAP_NIS | SW | SW1 |
| | | | | S1 | s22 | h64 | H150 | HOK73 | - | - | - |
| | | | | S1 | s22 | h64 | H149 | HOK74 | - | - | - |
| | | | | S1 | s22 | h64 | H151 | HOK75 | - | - | - |

| | | | | | | | | | | | |
|---|---|---|---|---|---|---|---|---|---|---|---|
| 50 | Minami, Sapporo, Hokkaido | 42.961 | 141.347 | S1 | s22 | h104 | H152 | - | - | - | - |
| | | | | S1 | s47 | h105 | H153 | - | - | - | - |
| | | | | S1 | s22 | h104 | H152 | - | - | - | - |
| | | | | S1 | s22 | h64 | H154 | - | s50_HOK_SAP_MIN | SW | SW1 |
| 51 | Minamishimamatsu, Eniwa, Hokkaido | 42.896 | 141.588 | S1 | s22 | h106 | H155 | - | s51_HOK_ENI | SW | SW1 |
| | | | | S1 | s22 | h64 | H156 | HOK76 | - | - | - |
| | | | | S1 | s22 | h64 | H157 | HOK77 | - | - | - |
| | | | | S1 | s45 | h107 | H158 | HOK78 | - | - | - |
| 52 | Kasuga, Chitose, Hokkaido | 42.820 | 141.638 | S1 | s22 | h64 | H157 | - | s52_HOK_CHI | SW | SW1 |
| | | | | S1 | s48 | h108 | H159 | - | - | - | - |
| | | | | S1 | s22 | h64 | H97 | - | - | - | - |
| | | | | S1 | s49 | h109 | H160 | - | - | - | - |
| 53 | Abira, Hokkaido | 42.824 | 141.834 | S1 | s50 | h110 | H161 | - | s53_HOK_ABI2 | SW | Admixed |
| | | | | S1 | s34 | h78 | H162 | - | - | - | - |
| | | | | S1 | s51 | h111 | H163 | - | - | - | - |
| | | | | S1 | s34 | h78 | H164 | - | s53_HOK_ABI1 | SW | Admixed |
| 54 | Miyamae, Tomakomai, Hokkaido | 42.607 | 141.484 | S1 | s22 | h64 | H165 | - | - | - | - |
| | | | | S1 | s22 | h64 | H165 | - | - | - | - |
| 55 | Shiraoi, Hokkaido | 42.541 | 141.331 | S1 | s34 | h78 | H166 | HOK79 | - | - | - |
| | | | | S1 | s52 | h112 | H167 | HOK80 | - | - | - |
| | | | | S1 | s52 | h112 | H167 | HOK81 | - | - | - |
| 56 | Niki, Hokkaido | 43.136 | 140.749 | S1 | s22 | h64 | H168 | - | s56_HOK_NIK2 | SW | SW1 |
| | | | | S1 | s53 | h113 | H169 | - | s56_HOK_NIK1 | SW | SW1 |
| | | | | S1 | s22 | h64 | H168 | - | - | - | - |
| | | | | S1 | s54 | h114 | H170 | - | - | - | - |
| 57 | Otani, Rankoshi, Hokkaido | 42.796 | 140.497 | S1 | s55 | h115 | H171 | HOK82 | - | - | - |
| | | | | S1 | s56 | h116 | H172 | - | s57_HOK_RAN1 | SW | Admixed |
| | | | | S1 | s55 | h117 | H173 | HOK83 | s57_HOK_RAN2 | SW | Admixed |
| | | | | S1 | s55 | h117 | H174 | HOK84 | - | - | - |
| 58 | Kombu, Rankoshi, Hokkaido | 42.780 | 140.601 | S1 | s57 | h118 | H175 | - | - | - | - |
| | | | | S1 | s55 | h117 | H173 | - | - | - | - |
| | | | | S1 | s55 | h117 | H173 | - | - | - | - |
| 59 | Nishisekinai, Date, Hokkaido | 42.487 | 140.874 | S1 | s22 | h64 | H176 | HOK85 | s59_HOK_DAT1 | SW | SW2 |
| | | | | S1 | s58 | h119 | H177 | HOK86 | s59_HOK_DAT2 | SW | SW2 |
| | | | | S1 | s59 | h120 | H178 | HOK87 | - | - | - |
| | | | | S1 | s58 | h119 | H177 | HOK88 | - | - | - |
| 60 | Imakane, Hokkaido | 42.422 | 140.019 | S1 | s60 | h121 | H179 | - | s60_HOK_IMA | SW | SW2 |
| | | | | S1 | s60 | h121 | H180 | - | - | - | - |
| | | | | S1 | s60 | h121 | H179 | - | - | - | - |
| 61 | Yakumo, Hokkaido | 42.256 | 140.245 | S1 | s41 | h122 | H181 | HOK89 | s61_HOK_YAK | SW | SW2 |
| | | | | S1 | s41 | h123 | H182 | HOK90 | - | - | - |
| | | | | S1 | s41 | h122 | H181 | HOK91 | - | - | - |
| | | | | S1 | s41 | h123 | H182 | HOK92 | - | - | - |
| 62 | Hongo, Hokuto, Hokkaido | 41.891 | 140.634 | S1 | s61 | h124 | H183 | - | - | - | - |
| | | | | S1 | s61 | h124 | H183 | - | - | - | - |
| | | | | S1 | s61 | h124 | H183 | - | - | - | - |
| | | | | S1 | s61 | h124 | H183 | - | - | - | - |
| 63 | Marumori, Miyagi | 37.916 | 140.773 | S1 | s34 | - | - | - | - | - | - |
| | | | | S1 | s34 | - | - | - | - | - | - |
| | | | | S1 | s34 | - | - | - | - | - | - |

| | | | | | | | | | | | |
|---|---|---|---|---|---|---|---|---|---|---|---|
| 64 | Miyashiro, Fukushima, Fukushima | 37.812 | 140.486 | S1 | s34 | - | - | - | - | - | - |
| | | | | S1 | s34 | - | - | - | - | - | - |
| | | | | S1 | s34 | - | - | - | - | - | - |
| 65 | Oda, Fukushima, Fukushima | 37.694 | 140.418 | S1 | s50 | - | - | - | - | - | - |
| | | | | S1 | s34 | - | - | - | - | - | - |
| 66 | Yonezawa, Yamagata | 37.958 | 140.070 | S1 | s34 | - | - | - | - | - | - |
| | | | | S1 | s34 | - | - | - | - | - | - |
| | | | | S1 | s34 | - | - | - | - | - | - |
| 67 | Agano river, Niigata | 37.742 | 139.319 | S1 | s34 | - | - | - | - | - | - |
| | | | | S1 | s34 | - | - | - | - | - | - |
| | | | | S1 | s34 | - | - | - | - | - | - |
| 68 | Tanagura, Fukushima | 36.994 | 140.398 | S1 | s34 | - | - | - | - | - | - |
| | | | | S1 | s34 | - | - | - | - | - | - |
| | | | | S1 | s34 | - | - | - | - | - | - |
| 69 | Daigo, Ibaraki | 36.765 | 140.357 | S1 | s34 | - | - | - | - | - | - |
| 70 | Kaname river, Kanagawa | 35.359 | 139.299 | S1 | s34 | - | - | - | - | - | - |
| | | | | S1 | s34 | - | - | - | - | - | - |
| | | | | S1 | s34 | - | - | - | - | - | - |

**Table S2.** List of specimens, sampling locations, mitochondrial haplotypes, and GenBank accession numbers of *Barbatula* species in continental East Asia

| Site No. | Locality | Latitude | Longitude | mtDNA Haplotype | | nDNA *RAG1* OTU | SNPs OTU | SNPs Group | | Acces. No. | | | | | |
|---|---|---|---|---|---|---|---|---|---|---|---|---|---|---|---|
| | | | | *cytb* + 12S | All mtDNA regions | | | Lineage | Sublineage | mtDNA *cytb* | mtDNA *COI* | mtDNA D-loop | mtDNA 12S rRNA, tRNA-Val, and 16S rRNA | nDNA *RAG1* | SNPs |
| *Barbatula* sp. | | | | | | | | | | | | | | | |
| 71 | Amur river, Mogol, Russia | 52.127 | 140.375 | h125 | H184 | - | - | - | - | | | | | | |
| | | | | h125 | H185 | - | - | - | - | | | | | | |
| | | | | h126 | H186 | RUS1 | s71_RUS_MOG1 | C | C3 | | | | | | |
| | | | | h127 | H187 | - | s71_RUS_MOG2 | C | C3 | | | | | | |
| 72 | Amur river, Chadbah, Russia | 53.043 | 141.150 | h125 | H188 | RUS2 | - | - | - | | | | | | |
| | | | | h128 | H189 | - | s72_RUS_CHA | C | C3 | | | | | | |
| 73 | Amur river, Anikinskiye, Russia | 52.760 | 140.177 | h129 | H190 | RUS3 | s73_RUS_ANI1 | C | C3 | | | | | | |
| | | | | h130 | H191 | RUS4 | s73_RUS_ANI2 | C | C3 | | | | | | |
| 74 | Amur river, B.Hadya, Russia | 48.768 | 139.960 | h131 | H192 | - | s74_RUS_HAD | C | C2 | | | | | | |
| | | | | h131 | H192 | RUS5 | - | - | - | | | | | | |
| 75 | Maksimovka, Primorsky Krai, Russia | 46.114 | 137.898 | h132 | H193 | - | s75_RUS_MAK | B | B | | | | | | |
| | | | | h132 | H194 | RUS6 | - | - | - | | | | | | |
| 76 | Tayozhanya, Primorsky Krai, Russia | 45.471 | 136.689 | h133 | H195 | - | s76_RUS_TAY | B | B | | | | | | |
| | | | | h133 | H196 | - | - | - | - | | | | | | |
| 77 | Krai Bikin river, Primorsky Krai, Russia | 46.827 | 134.217 | h134 | H197 | - | s77_RUS_BIK | F | F | | | | | | |
| | | | | h135 | H198 | - | - | - | - | | | | | | |
| 78 | Arsenjevka, Primorsky Krai, Russia | 44.385 | 133.459 | h136 | H199 | - | - | - | - | | | | | | |
| | | | | - | - | - | s78_RUS_ARS | A | A2 | | | | | | |
| 79 | Vladivostok, Primorsky Krai, Russia | 43.212 | 132.069 | h137 | H200 | - | - | - | - | | | | | | |
| | | | | h137 | H201 | - | - | - | - | | | | | | |
| 80 | Shkotovsky, Primorsky Krai, Russia | 43.309 | 132.768 | h138 | H202 | - | s80_RUS_SHK | A | A1 | | | | | | |
| | | | | h139 | H203 | - | - | - | - | | | | | | |
| 81 | Fokino, Primorsky Krai, Russia | 42.962 | 132.418 | h140 | H204 | - | - | - | - | | | | | | |
| 82 | Novoneghino Sukhodal river, Primorsky Krai, Russia | 43.227 | 132.544 | h141 | H205 | - | s82_RUS_NOV | A | A1 | | | | | | |
| | | | | h142 | H206 | - | - | - | - | | | | | | |
| *Barbatula nuda* | | | | | | | | | | | | | | | |
| 83 | Benxi, Liaoning, China | 41.081 | 124.194 | h143 | H207 | - | s83_CHI_BEN | F | F | | | | | | |
| | | | | h144 | H208 | - | - | - | - | | | | | | |
| *Barbatula* cf. *toni* | | | | | | | | | | | | | | | |
| 84 | Hebei, China | 41.699 | 115.774 | h145 | H209 | - | - | - | - | | | | | | |
| | | | | h146 | H210 | - | s84_CHI_HEB | C | C1 | | | | | | |
| *Barbatula* sp. | | | | | | | | | | | | | | | |
| 85 | Goseong, Gangwon, Korea | 38.371 | 128.434 | h147 | H211 | - | s85_KOR_GOS | A | A1 | | | | | | |
| | | | | h147 | H211 | - | - | - | - | | | | | | |
| *Barbatula* aff. *nuda* | | | | | | | | | | | | | | | |
| 86 | Pyeongchang, Gangwon, Korea | 37.665 | 128.586 | h148 | H212 | KOR1 | s86_KOR_PYE | D | D | | | | | | |
| | | | | h148 | H213 | KOR2 | - | - | - | | | | | | |
| | | | | h149 | H214 | KOR3 | - | - | - | | | | | | |
| | | | | h149 | H214 | KOR4 | - | - | - | | | | | | |
| 87 | Samcheok, Gangwon, Korea | 37.412 | 129.101 | h150 | H215 | KOR5 | s87_KOR_SAM | D | D | | | | | | |
| | | | | h151 | H216 | KOR6 | - | - | - | | | | | | |

| | | | | | | | | | |
|---|---|---|---|---|---|---|---|---|---|
| 88 | Hwacheon, Gangwon, Korea | 38.061 | 127.513 | h152 | H217 | KOR7 | s88_KOR_HWA | E | E |
| | | | | h153 | H218 | KOR8 | - | - | - |
| 89 | Inje, Gangwon, Korea | 38.083 | 128.186 | h152 | H219 | KOR9 | - | - | - |
| | | | | h152 | H219 | KOR10 | - | - | - |
| 90 | Weonju, Gangwon-do, Korea | 37.220 | 128.086 | h154 | H220 | KOR11 | s90_KOR_WEO | E | E |
| | | | | h155 | H221 | KOR12 | - | - | - |

**Table S3.** List of specimens and GenBank accesssion numbers of outgroup

| Family | Species | Site name | Acces. No. | | | | | | References |
|---|---|---|---|---|---|---|---|---|---|
| | | | complete | mtDNA *cytb* | mtDNA *COI* | mtDNA D-loop | mtDNA 12S rRNA, tRNA-Val, and 16S | nDNA *RAG1* | |
| Nemacheilidae | *Triplophysa sellaefer* | Lvliang, Shanxi, China | - | | | | | - | This study |
| Nemacheilidae | *Triplophysa robusta* | Dingxi, Gansu, China | - | | | | | - | This study |
| Nemacheilidae | *Triplophysa luochengensis* | Pearl, Guangxi, China | - | | | | | PP315886 | Šlechtová et al. (2024) |
| Nemacheilidae | *Triplophysa labiata* | Lake Balkash, Almaty, Kazachstan | - | | | | | PP315842 | Šlechtová et al. (2024) |
| Nemacheilidae | *Lefua costata* | Pyeongchang, Gangwon, Korea | - | | | | | - | This study |
| Nemacheilidae | *Lefua echigonia* | Ohtawara, Tochigi, Japan | - | | | | | - | This study |
| Nemacheilidae | *Lefua nikkonis* | - | NC_027662 | - | - | - | - | - | Miya et al. (2015) |
| Cobitidae | *Cobitis biwae* | Matsumoto, Nagano, Japan | - | | | | | - | This study |
| Cobitidae | *Misgurnus anguillicaudatus* | Hirosaki, Aomori, Japan | - | | | | | - | This study |
| Cobitidae | *Kichulchoia multifasciata* | - | AP011337 | - | - | - | - | - | Unpublished |
| Cobitidae | *Cobitis elongatoides* | - | NC_023947 | - | - | - | - | - | Huang et al. (2014) |
| Cobitidae | *Koreocobitis rotundicaudata* | - | NC_018755 | - | - | - | - | - | Kim and Bang (2012) |


Huang, S., Tomljanovic, T., Tian, X., Wang, Y., Cao, X., 2016. The complete mitochondrial genome of natural *Cobitis elongatoides* (Cypriniformes: Cobitidae). *Mitochondrial DNA Part A* 27, 189–190. https://doi.org/10.3109/19401736.2013.879654.

Kim, K.Y., Bang, I.C., 2012. Phylogeny and speciation time estimation of two *Koreocobitis* species (Teleostei; Cypriniformes; Cobitidae) endemic to Korea inferred from their complete mitogenomic sequences. *Genes Genom.* 34, 35–42. https://doi.org/10.1007/s13258-011-0139-5.

Miya, M., Sato, Y., Fukunaga, T., Sado, T., Poulsen, J.Y., Sato, K., Minamoto, T., Yamamoto, S., Yamanaka, H., Araki, H., Kondoh, M., Iwasaki, W., 2015. MiFish, a set of universal PCR primers for metabarcoding environmental DNA from fishes: detection of more than 230 subtropical marine species. *R. Soc. Open Sci.* 2, 150088. https://doi.org/10.1098/rsos.150088.

Šlechtová, V., Dvořák, T., Freyhof, J., Kottelat, M., Levin, B., Golubtsov, A., Šlechta, V., Bohlen, J., 2024. Reconstructing the evolutionary history of freshwater fishes (Nemacheilidae) across Eurasia since early Eocene. *eLife* 13, e101080. https://doi.org/10.7554/eLife.101080.1.

**Table S4.** Usage primers and PCR conditions

| Region | Primer | Sequence 5'–3' | Reference | Annealing temperature |
|---|---|---|---|---|
| *cytb* | L15285 | CCCTAACCCGVTTCTTYGC | Inoue et al. (2000) | 50 °C |
| | H15915 | ACCTCCGATCTYCGGATTACAAGAC | Aoyama et al. (2000) | |
| *COI* | Fish-F1 | TCAACCAACCACAAAGACATTGGCAC | Ward et al. (2005) | 52 °C |
| | Fish-R1 | TAGACTTCTGGGTGGCCAAAGAATCA | Ward et al. (2005) | |
| D-loop | Pro S | GCATCGGTCTTGTAATCCGAAGAT | Sakai et al. (2003) | 62 °C |
| | Phe AS | GGACCAAGCCTTTGTGCATGCGGAG | Sakai et al. (2003) | |
| 12S rRNA, tRNA-Val and 16S rRNA | L708-12S | TTAYACATGCAAGTMTCCGC | Miya et al. (2004) | 54 °C |
| | H1471-16S | CACCAAGTTCGGTAGGTTTAT | Wang et al. (2016) | |
| RAG1 | RAG1F | AGCTGTAGTCAGTAYCACAARATG | Quenouille et al. (2004) | 54 °C |
| | RAG-RV1 | TCCTGRAAGATYTTGTAGAA | Šlechtová et al. (2007) | |


Aoyama, J., Watanabe, S., Ishikawa, S., Nishida, M., Tsukamoto, K., 2000. Are morphological characters distinctive enough to discriminate between two species of freshwater eels, *Anguilla celebesensis* and *A. interioris*? *Ichthyol. Res.* 47, 157–161. https://doi.org/10.1007/BF02684236.

Inoue, G.J., Miya, M., Tsukamoto, K., Nishida, M., 2000. Complete mitochondrial DNA sequence of the Japanese sardine *Sardinops melanostictus*. *Fish. Sci.* 66, 924–932. https://doi.org/10.1046/j.1444-2906.2000.00148.x.

Miya, M., Satoh, T.R., Nishida, M., 2005. The phylogenetic position of toadfishes (order Batrachoidiformes) in the higher ray-finned fish as inferred from partitioned Bayesian analysis of 102 whole mitochondrial genome sequences. *Biol. J. Linn. Soc.* 85, 289–306. https://doi.org/10.1111/j.1095-8312.2005.00483.

Quenouille, B., Bermingham, E., Planes, S., 2004. Molecular systematics of the damselfishes (Teleostei: Pomacentridae): Bayesian phylogenetic analyses of mitochondrial and nuclear DNA sequences. *Mol. Phylogenet. Evol.* 31, 66–88. https://doi.org/10.1016/S1055-7903(03)00278-1.

Sakai, T., Mihara, M., Shitara, H., Yonekawa, H., Hosoya, K., Miyazaki, J., 2003. Phylogenetic relationships and intraspecific variations of loaches of the genus *Lefua* (Balitoridae, Cypriniformes). *Zool. Sci.* 20, 501–514. https://doi.org/10.2108/zsj.20.501.

Šlechtová, V., Bohlen, J., Tan, H.H., 2007. Families of Cobitoidea (Teleostei; Cypriniformes) as revealed from nuclear genetic data and the position of the mysterious genera *Barbucca*, *Psilorhynchus*, *Serpenticobitis* and *Vaillantella*. *Mol. Phylogenet. Evol.* 44, 1358–1365. https://doi.org/10.1016/j.ympev.2007.02.019.

Wang, Y., Shen, Y., Feng, C., Zhao, K., Song, Z., Zhang, Y., Yang, L., He, S., 2016. Mitogenomic perspectives on the origin of Tibetan loaches and their adaptation to high altitude. *Sci. Rep.* 6, 29690. https://doi.org/10.1038/srep29690.

Ward, R.D., Zemlak, T.S., Innes, B.H., Last, P.R., Hebert, P.D.N., 2005. DNA barcoding Australia's fish species. *Philos. Trans. R. Soc. B* 360, 1847–1857. https://doi.org/10.1098/rstb.2005.1716.

**Table S5.** Details about SNPs filtering

| . | Usage samples | Filtering details | Number of SNPs | Analysis |
|---|---|---|---|---|
| *de novo* dataset | | | | |
| *de novo* dataset 1 | All (133 samples) | SNPs shared by all samples were retained (-R 1.0), singleton SNPs were removed (--min-mac 2), maximum observed heterozygosity was 0.5 (--max-obs-het 0.5), and one SNP per locus was selected (--write-single-snp) using the *populations* program in Stacks ver. 2.66 | 3,097 SNPs | Neighbor-net |
| *de novo* dataset 2 | All | From *de novo* dataset 1, SNPs with linkage disequilibrium (LD) above 0.8 were excluded using PLINK ver. 1.90 (--indep-pairwise 100 5 0.8) | 2,178 SNPs | ADMIXTURE |
| *de novo* dataset 3 | *B. oreas* only (114 samples) | The same procedure as for *de novo* dataset 1 was applied | 8,178 SNPs | Neighbor-net |
| *de novo* dataset 4 | *B. oreas* only | From *de novo* dataset 3, SNPs with linkage disequilibrium (LD) above 0.8 were excluded using PLINK | 7,773 SNPs | ADMIXTURE |
| *de novo* dataset 5 | Russia, China, and Korea (19 samples) | The same procedure as for *de novo* dataset 1 was applied, and SNPs with linkage disequilibrium (LD) above 0.8 were excluded using PLINK | 2,116 SNPs | ADMIXTURE |
| mapping dataset | | | | |
| mapping dataset 1 | All | Indels were removed (--remove-indels), and SNPs were filtered using VCFtools v.0.1.16 to retain sites with quality ≥ 30 (--minQ 30), genotype quality ≥ 20 (--minGQ 20), depth ≥ 8 (--minDP 8), mean depth between 10 and 300 (--min-meanDP 10, --max-meanDP 300), biallelic sites (--min-alleles 2, --max-alleles 2), minor allele frequency ≥ 0.01 (--maf 0.01), and missing rate ≤ 20% (--max-missing 0.8) | 159,567 SNPs | SVDquartets<br>Neighbor-net |
| mapping dataset 2 | All | From mapping dataset 1, invariant sites were removed | 132,537 SNPs | IQ-TREE |
| mapping dataset 3 | All | From mapping dataset 1, SNPs with linkage disequilibrium (LD) above 0.1 were excluded using PLINK (--indep-pairwise 100 5 0.1) | 19,243 SNPs | ADMIXTURE |
| mapping dataset 4 | All | From mapping dataset 3, outlier SNPs were excluded using pcadapt ver. 4.4.0 (K = 5, min.maf = 0.05, α = 0.1), and SNPs were thinned to a minimum distance of 2,000 bp using VCFtools (--thin 2000) | 8,178 SNPs | *f*-branch |
| mapping dataset 5 | All | From mapping dataset 1, SNPs shared by all samples were retained using VCFtools | 30,314 SNPs | TreeMix |
| mapping dataset 6 | *B. oreas* only | The same procedure as for mapping dataset 1 was applied | 111,035 SNPs | Neighbor-net |
| mapping dataset 7 | *B. oreas* only | From mapping dataset 6, SNPs with linkage disequilibrium (LD) above 0.1 were excluded using PLINK | 13,125 SNPs | ADMIXTURE<br>DAPC<br>PCA<br>pairwise $F_{ST}$<br>AMOVA |

| | | | | |
|---|---|---|---|---|
| mapping dataset 8 | *B. oreas* only | From mapping dataset 7, outlier SNPs were excluded using pcadapt, and SNPs were thinned to a minimum distance of 2,000 bp using VCFtools | 8,296 SNPs | divMigrate |
| mapping dataset 9 | Russia, China, and Korea | The same procedure as for mapping dataset 1 was applied, except that singleton SNPs were removed (--min-mac 2) instead of applying a minor allele frequency filter (MAF > 0.01), and SNPs with linkage disequilibrium (LD) above 0.3 were excluded using PLINK | 7,247 SNPs | ADMIXTURE |
| mapping dataset 10 | Lineages A–C only (13 samples) | The same procedure as for mapping dataset 9 was applied | 2,956 SNPs | PCA<br>pairwise $F_{ST}$ |

**Table S6.** Genetic diversity of *Barbatula oreas* based on the mtDNA D-loop, *COI*, and 12S rRNA regions

| | N | D-loop | | | *COI* | | | 12S rRNA | | |
|---|---|---|---|---|---|---|---|---|---|---|
| | | Nh | Hd | π | Nh | Hd | π | Nh | Hd | π |
| Northern Clade | | | | | | | | | | |
| N1 | 68 | 41 | 0.967 | 0.006 | 15 | 0.539 | 0.002 | 16 | 0.785 | 0.0015 |
| N2 | 12 | 8 | 0.939 | 0.004 | 3 | 0.667 | 0.001 | 2 | 0.303 | 0.0004 |
| Subtotal | 80 | 49 | 0.975 | 0.010 | 18 | 0.661 | 0.003 | 17 | 0.752 | 0.0014 |
| Southern Clade | | | | | | | | | | |
| S1 | 104 | 61 | 0.982 | 0.006 | 22 | 0.872 | 0.004 | 19 | 0.437 | 0.0008 |
| S2 | 26 | 16 | 0.951 | 0.004 | 6 | 0.465 | 0.002 | 5 | 0.289 | 0.0005 |
| S3 | 8 | 5 | 0.786 | 0.002 | 3 | 0.679 | 0.001 | 1 | - | - |
| S4 | 20 | 13 | 0.932 | 0.003 | 6 | 0.579 | 0.002 | 6 | 0.658 | 0.0011 |
| Subtotal | 158 | 94 | 0.989 | 0.011 | 37 | 0.923 | 0.013 | 30 | 0.681 | 0.0016 |
| Total | 238 | 143 | 0.992 | 0.019 | 55 | 0.928 | 0.022 | 47 | 0.832 | 0.0031 |

N: number of examined specimens, Nh: number of haplotypes, Hd: haplotype diversity, π: nucleotide diversity.

**Table S7.** Pairwise $F_{ST}$ values (a) between the two major clades and (b) among the six subclades based on the mtDNA *cytb* region

(a)

| | Northern Clade | Southern Clade |
|---|---|---|
| Northern Clade | | |
| Southern Clade | 0.691 | |

(b)

| | N1 | N2 | S1 | S2 | S3 | S4 |
|---|---|---|---|---|---|---|
| N1 | | | | | | |
| N2 | 0.752 | | | | | |
| S1 | 0.883 | 0.907 | | | | |
| S2 | 0.892 | 0.922 | 0.863 | | | |
| S3 | 0.898 | 0.920 | 0.839 | 0.872 | | |
| S4 | 0.864 | 0.905 | 0.783 | 0.799 | 0.857 | |

**Table S8.** Genetic distances (*p* -distance) among species of the genus *Barbatula* based on the mtDNA *cytb* region

| | 1 | 2 | 3 | 4 | 5 | 6 | 7 | 8 |
|---|---|---|---|---|---|---|---|---|
| 1. *Barbatula oreas* (Hokkaido & Sakhalin) | | | | | | | | |
| 2. *B.* sp. (Maksimovka, Primorsky Krai, Russia) | 0.066 | | | | | | | |
| 3. *B.* sp.1* (Arsenjevka, Primorsky Krai, Russia) | 0.071 | 0.063 | | | | | | |
| 4. *B. nuda* (Liaoning, China) | 0.056 | 0.066 | 0.064 | | | | | |
| 5. *B.* cf. *toni* (Hebei, China) | 0.079 | 0.044 | 0.064 | 0.078 | | | | |
| 6. *B.* sp.3* (Goseong, Gangwon, Korea) | 0.056 | 0.067 | 0.058 | 0.030 | 0.077 | | | |
| 7. *B.* aff. *nuda* (Western Korea) | 0.151 | 0.147 | 0.142 | 0.156 | 0.154 | 0.144 | | |
| 8. *B.* aff. *nuda* (Eastern Korea) | 0.118 | 0.103 | 0.105 | 0.120 | 0.105 | 0.114 | 0.157 | |

*Complying with Chen et al. (2019)

**Table S9.** Estimated divergence times (Ma) and 95% highest posterior density (HPD) intervals at major nodes of the time-calibrated phylogenetic tree, based on (1) molecular clocks and (2) fossil calibration points

| Node | (1) Molecular clock | | (2) Calibration | |
|---|---|---|---|---|
| Subclades of Northern Clade | 0.58 | [0.34–0.85] | 1.04 | [0.57–1.60] |
| Subclades of Southern Clade | 0.83 | [0.54–1.15] | 1.41 | [0.82–2.04] |
| Northern Clade and Southern Clade | 1.35 | [0.90–1.83] | 2.32 | [1.40–3.33] |
| *Barbatula* species | 5.90 | [3.63–8.67] | 10.57 | [6.61–14.85] |

**Table S10.** Cross-validation errors from ADMIXTURE analyses for *de novo* and mapping SNP datasets.

(a) *de novo* dataset 2 (all samples)

| K | Cross-validation error |
|---|---|
| 2 | 0.221 |
| 3 | 0.199 |
| 4 | 0.187 |
| 5 | 0.185 |
| 6 | 0.186 |
| 7 | 0.177 |
| 8 | 0.175 |
| 9 | 0.181 |
| 10 | 0.175 |

(b) *de novo* dataset 4 (*B. oreas* only)

| K | Cross-validation error |
|---|---|
| 2 | 0.209 |
| 3 | 0.206 |
| 4 | 0.227 |
| 5 | 0.236 |
| 6 | 0.224 |
| 7 | 0.204 |
| 8 | 0.203 |
| 9 | 0.192 |
| 10 | 0.192 |
| 11 | 0.194 |
| 12 | 0.204 |

(c) *de novo* dataset 5 (continental)

| K | Cross-validation error |
|---|---|
| 2 | 1.135 |
| 3 | 1.382 |
| 4 | 1.222 |
| 5 | 1.683 |
| 6 | 1.375 |

(d) mapping dataset 3 (all samples)

| K | Cross-validation error |
|---|---|
| 2 | 0.298 |
| 3 | 0.281 |
| 4 | 0.271 |
| 5 | 0.267 |
| 6 | 0.257 |
| 7 | 0.261 |
| 8 | 0.278 |
| 9 | 0.269 |
| 10 | 0.283 |

(e) mapping dataset 7 (*B. oreas* only)

| K | Cross-validation error |
|---|---|
| 2 | 0.325 |
| 3 | 0.305 |
| 4 | 0.302 |
| 5 | 0.294 |
| 6 | 0.300 |
| 7 | 0.310 |
| 8 | 0.316 |
| 9 | 0.315 |
| 10 | 0.336 |
| 11 | 0.359 |
| 12 | 0.372 |

(f) mapping dataset 9 (continental)

| K | Cross-validation error |
|---|---|
| 2 | 1.272 |
| 3 | 1.491 |
| 4 | 1.624 |
| 5 | 1.741 |
| 6 | 1.797 |

**Table S11.** Pairwise $F_{ST}$ (below the diagonal) and p-values (above the diagonal): (a) among the three major lineages and (b) among the six sublineages based on 2,956 nuclear SNPs

(a)

| | A | B | C |
|---|---|---|---|
| A | | 0.067 | 0.004 |
| B | 0.390 | | 0.029 |
| C | 0.313 | 0.373 | |

(b)

| | A1 | A2 | B | C1 | C2 | C3 |
|---|---|---|---|---|---|---|
| A1 | | 1.000 | 0.111 | 1.000 | 1.000 | 0.016 |
| A2 | 0.528 | | 0.325 | 1.000 | 1.000 | 0.174 |
| B | 0.493 | 0.683 | | 1.000 | 1.000 | 0.048 |
| C1 | 0.556 | 0.778 | 0.681 | | 1.000 | 0.168 |
| C2 | 0.579 | 0.774 | 0.657 | 0.755 | | 0.172 |
| C3 | 0.447 | 0.543 | 0.487 | 0.485 | 0.369 | |

**Table S12.** Directional relative migration rates among the nine subclusters inferred using divMigrate

| Populations | NM1 | NM2 | NW | MW | SW1 | SW2 | SE1 | SE2 | NE |
|---|---|---|---|---|---|---|---|---|---|
| NM1 | NA | **0.641** | **1.000** | **0.306** | **0.077** | 0.039 | 0.022 | 0.013 | **0.088** |
| NM2 | 0.152 | NA | **0.270** | **0.106** | 0.046 | 0.026 | 0.017 | 0.010 | 0.050 |
| NW | 0.055 | 0.069 | NA | 0.093 | 0.052 | 0.031 | 0.018 | 0.011 | 0.034 |
| MW | 0.027 | 0.033 | 0.096 | NA | 0.047 | 0.030 | 0.015 | 0.010 | 0.027 |
| SW1 | 0.035 | 0.046 | **0.220** | **0.184** | NA | 0.166 | 0.036 | 0.019 | 0.069 |
| SW2 | 0.036 | **0.048** | **0.261** | **0.198** | **0.518** | NA | 0.037 | 0.018 | 0.072 |
| SE1 | **0.034** | **0.047** | **0.242** | **0.149** | **0.123** | **0.066** | NA | 0.068 | **0.145** |
| SE2 | **0.031** | **0.043** | **0.250** | **0.126** | **0.116** | **0.060** | **0.281** | NA | **0.125** |
| NE | 0.044 | 0.064 | **0.234** | **0.133** | 0.087 | 0.052 | 0.052 | 0.024 | NA |

This table shows migration rates from column populations to row populations. Bold text indicates unidirectional migration supported by 1,000 bootstrap replicates
